# KCNQ5 controls perivascular adipose tissue-mediated vasodilation

**DOI:** 10.1101/2023.07.26.550617

**Authors:** Dmitry Tsvetkov, Johanna Schleifenbaum, Yibin Wang, Mario Kassmann, Maya M Polovitskaya, Sebastian Schütze, Michael Rothe, Friedrich C Luft, Thomas J Jentsch, Maik Gollasch

## Abstract

**Background:** Small arteries exhibit resting tone, a partially contracted state that maintains arterial blood pressure. In arterial smooth muscle cells (SMCs), potassium channels control contraction and relaxation. Perivascular adipose tissue (PVAT) has been shown to exert anticontractile effects on the blood vessels. However, the mechanisms by which PVAT signals small arteries, and their relevance, remain largely unknown. We aimed to uncover key molecular components in adipose-vascular coupling.

**Methods:** A wide-spectrum of genetic mouse models targeting *Kcnq3, Kcnq4* and *Kcnq5* genes (*Kcnq3*^−/−^, *Kcnq4*^−/−^, *Kcnq5*^−/−^, *Kcnq5*^dn/dn^, *Kcnq4*^−/−^*/Kcnq5*^dn/dn^, *Kcnq4*^−/−^*/Kcnq5*^−/−^), telemetry blood pressure measurements, targeted lipidomics, and RNA-Seq profiling, wire-myography, patch-clamp, and sharp-electrode membrane potential measurements were used.

**Results:** We show that PVAT causes SMC KCNQ5 (K_V_7.5) channels to hyperpolarize the membrane potential. This effect relaxes small arteries and regulates blood pressure. Oxygenation of polyunsaturated fats generates oxylipins, a superclass of lipid mediators. We identified numerous oxylipins released by PVAT that potentiate vasodilatory action in small arteries by opening SMC KCNQ5 channels.

**Conclusions:** Our results reveal a key molecular function of KCNQ5 channels in adipose-vascular coupling, translating PVAT signals, particularly oxylipins, to the central physiological function of vasoregulation. This novel pathway opens new therapeutic perspectives.

## Introduction

Hypertension is the leading cause of cardiovascular disease and premature death worldwide. ^1 2^ Maintained blood-pressure elevation involves increased peripheral vascular resistance. ^3^ Arterial vessels exhibit a resting tone that is influenced in part by perivascular adipose tissue (PVAT). The PVAT-vessel interaction has been termed adipose-vascular coupling and represents a fundamental physiological process of vascular tone regulation.^4 5^ In health, this coupling is beneficial; in disease and aging, perhaps not so^5^. However, how the PVAT signals are transmitted is not completely understood. Potassium channels are the main molecular components in smooth muscle cells (SMCs) mediating vascular relaxation by PVAT.^4^ We focused on potassium channels encoded by the *KCNQ* gene family, which are widely expressed in arterial smooth muscle cells (SMCs).^6^ Evidence exists suggesting that KCNQ potassium channels could serve as physiological intermediates in vasodilatory signaling.^7 8^ In that regard, the *KCNQ* gene family is remarkable. Mutations in all five *KCNQ* genes underlie diseases in humans including long QT (*KCNQ1*) syndrome^9^, epileptic encephalopathy (*KCNQ2/3*)^10 11^, deafness (*KCNQ4*)^12^, and complex central nervous system (CNS) symptoms including intellectual disability (ID) and epileptic encephalopathy (*KCNQ5*). ^13 14^ *Kcnq* mouse models generally display phenotypes resembling those of respective human diseases.^15 16 17^ Unsurprisingly, however, *Kcnq5*^−/−^ and *Kcnq*5^dn^ mice do not show the severe neurological phenotypes observed in patients with *de novo KCNQ5* missense mutations, which entail specific changes in biophysical channel properties.^18 19 13 14^ Since KCNQ2 channels are not expressed in the vasculature^6 20^ and because KCNQ1 channels are unlikely to be involved^21^, we focused on vasoregulation by potassium channels encoded by the *KCNQ3* (K_V_7.3), *KCNQ4* (K_V_7.4) and *KCNQ5* (K_V_7.5) genes. Using a wide-spectrum of genetic mouse models targeting *Kcnq3, Kcnq4* and *Kcnq5* genes (*Kcnq3*^−/−^, *Kcnq4*^−/−^, *Kcnq5*^−/−^, *Kcnq5*^dn/dn^, *Kcnq4*^−/−^*/Kcnq5*^dn/dn^, *Kcnq4*^−/−^*/Kcnq5*^−/−^) and targeted lipidomics profiling, we demonstrate that KCNQ5 in SMCs is a key molecular component in adipose-vascular coupling. KCNQ5 translates PVAT signals, particularly oxylipins, into vasoregulation and blood pressure control. Our results uncover key molecular components in adipose-vascular coupling and highlight the therapeutic potential of targeting PVAT against cardiovascular complications.

## Results

### KCNQ4/KCNQ5 channels promote vasodilation

Synthetic KCNQ2-5 (K_V_7.2-5) channel activators, such as flupirtine and retigabine, relax blood vessels of different organs and species.^22 23 24^ However, the target specificity of the effects is unclear.^25 26^ Wire myography was used to measure vasocontractions of mesenteric artery rings. PVAT was removed from the arterial rings ((–) PVAT). ^4 27 28^ We first studied the effects of flupirtine and retigabine in *Kcnq* mutant mice. *Kcnq4*^−/−^, *Kcnq5*^−/−^, and *Kcnq4*^−/−^/*Kcnq5*^−/−^ arteries exhibited attenuated relaxation in response to either KCNQ2–5 (K_V_7.2–5) channel activator, whereas arterioles from wild-type and *Kcnq3*^−/−^ dilated robustly (Fig. 1a,b,c; Fig. S1, Fig. S2). Targeted deletion of KCNQ4 and/or KCNQ5 had no effect on contraction induced by stimulation of alpha-1 adrenergic receptors, thromboxane A2 receptors (U46619), and arginine vasopressin (AVP) receptors (Fig. S3). The results suggest that deletion of KCNQ4/KCNQ5 impairs relaxation only upon pharmacological activation of the channels but has no role in vascular contraction induced by the humoral vasoconstrictors. Our results argue against an important role of SMCs KCNQ3 channels in vasoregulation. Constitutive gene disruption may entail secondary compensatory changes in gene expression.^29^ To exclude such changes in the non-targeted KCNQ channels and other vasoactive pathways in the mouse models used, we performed targeted bulk RNA-sequencing of the isolated arteries. Our results argue against significant alteration of mRNA levels of non-targeted KCNQ channels and other essential vasoregulatory targets (Fig. S4).

**Fig. 1.**
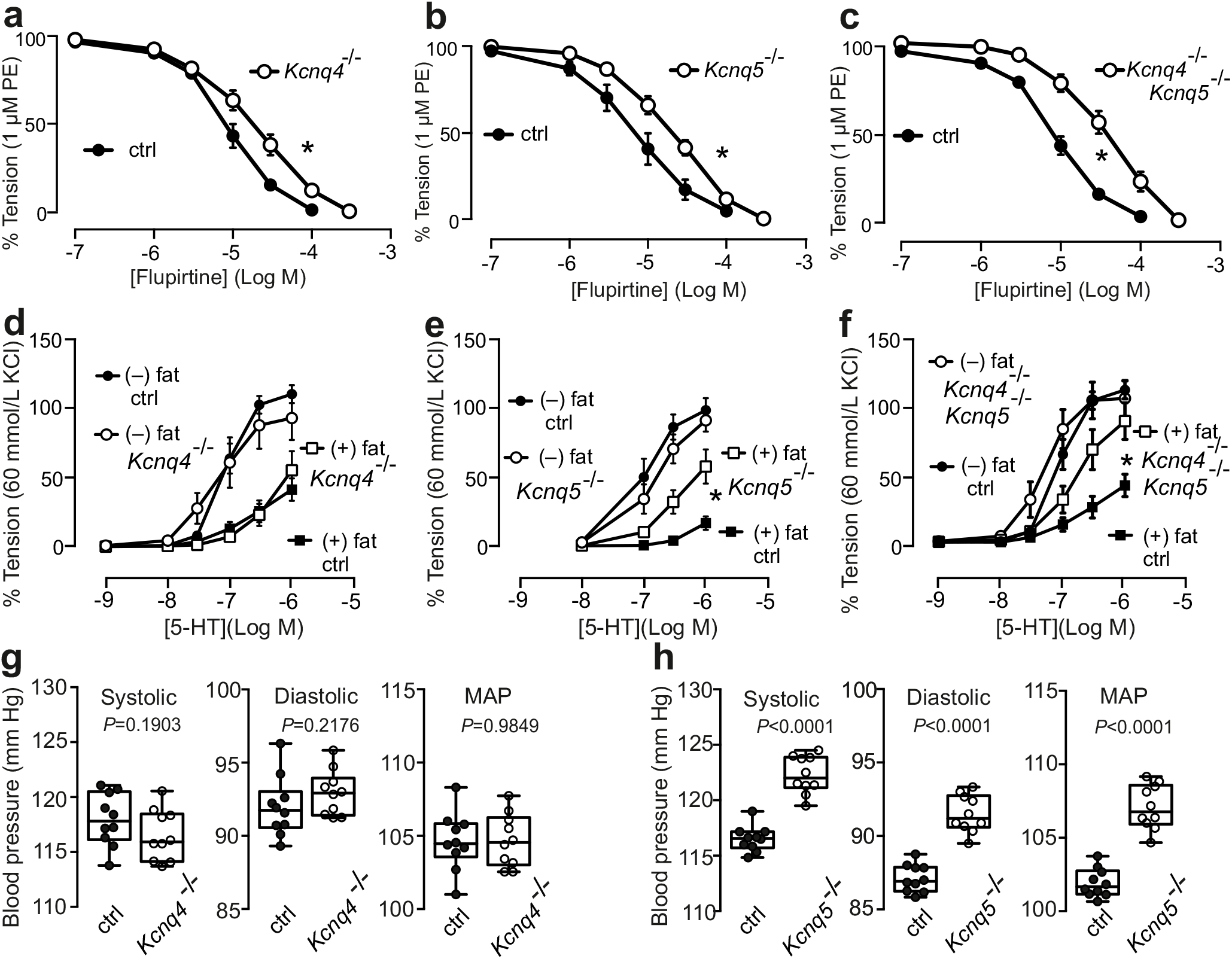
KCNQ5 channel is required for the arterial tone by PVAT and blood pressure control. Relaxation of (−) PVAT mesenteric arteries by flupirtine in *Kcnq4*^−/−^ (**a**), *Kcnq5*^−/−^ (**b**), *Kcnq4*^−/−^/*Kcnq5*^−/−^ or respective control (*Kcnq4*^+/+^, *Kcnq5*^+/+^, *Kcnq4*^+/+^/*Kcnq5*^+/+^) mice (**c**). Tension is expressed as a percentage of alpha1-adrenoreceptor agonist-induced contraction. Concentration-response relationships for serotonin (5-HT) induced contractions in (+) PVAT or (–) PVAT artery rings isolated from *Kcnq4*^−/−^ (**d**), *Kcnq5*^−/−^ (**e**), *Kcnq4*^−/−^*/Kcnq5*^−/−^ (**f**) and respective controls (*Kcnq4*^+/+^; *Kcnq5*^+/+^; *Kcnq4*^+/+^/*Kcnq5*^+/+^) mice. Tension is expressed as a percentage of 60 mM KCl-induced contractions. n≥6 arteries from N≥3 mice. Data are mean ± s.e.m. *P as determined by two-way ANOVA with Šidák post hoc test. Systolic, diastolic and mean arterial blood pressures (MAP) over 10 days in control (ctrl) *Kcnq4*^+/+^ (N=14) or *Kcnq4*^−/−^ mice (N=13) (**g**) *Kcnq5*^+/+^ or *Kcnq5*^−/−^ (N=8 mice per group) (**h**). Box plots show median and IQR, whiskers are max and min values. *P* as determined by two-tailed Mann–Whitney *U* test.

### KCNQ channels function independently of cAMP– and NO/cGMP–pathways

The role of the endothelium in the regulation of vascular tone is well established. Nitric oxide (NO), cGMP/protein kinase G and cAMP/protein kinase A pathways are involved in endothelial vasodilation elicited by a wide variety of stimuli, including hypoxia and hypercapnia and agonist/receptor interactions. In addition, large component of acetylcholine relaxation of mesenteric arteries is mediated by endothelial TRPV4 Ca^2+^ entry coupled to intermediate (IK)- and small (SK)-conductance Ca^2+^-sensitive potassium channels.^30^ Forskolin, which stimulates adenylyl cyclase to produce cAMP, induced similar relaxations in *Kcnq4*^−/−^, *Kcnq5*^−/−^, *Kcnq5*^dn/dn^, *Kcnq4*^−/−^*/Kcnq5*^dn/dn^, and respective wild-type mesenteric arteries (Fig. S5 a,b,c,d). Similarly, NO/cGMP– and TRPV4–IK/SK–dependent relaxations upon acetylcholine administration were similar in *Kcnq4*^−/−^, *Kcnq5*^−/−^, *Kcnq5*^dn/dn^, and respective wild-type arteries (Fig. S5 e,f,g). These results indicate that KCNQs channels are not involved in cAMP and NO/cGMP vasodilatory pathways, at least in mesenteric arteries. This is in line with our previous data demonstrating that PVAT control of arterial tone does not involve the endothelium or key potassium channels (BK_Ca_, SK, IK, K_ir_) controlled by endothelial factors *via* cAMP– and NO/cGMP–pathways (e.g. NO, endothelial-derived hyperpolarization factor (EDHF), potassium ions).^28^

### KCNQ5 channels regulate PVAT–mediated arterial tone and blood pressure

We next investigated the role of KCNQ channels in PVAT-mediated control of vascular tone. We hypothesized that modulation of KCNQ4 and/or KCNQ5 channel activity couples perivascular adipose tissue to vasodilation. Wire myography was used to measure vasocontractions of mesenteric artery rings. PVAT was either left intact (+) or removed (–) from the arterial rings. ^4 31^ *Kcnq4*^−/−^ mice exhibited normal anti-contractile effects of PVAT on arterial tone (Fig. 1d). In contrast, these anti-contractile effects of PVAT were attenuated in *Kcnq5*^−/−^ mice (Fig. 1e; Fig. S6). Similar results were observed in *Kcnq4*^−/−^/*Kcnq5*^−/−^ double knockout arteries (Fig. 1f). Since KCNQ activation may hyperpolarize the SMC membrane potential, we simultaneously measured membrane potential and force upon KCNQ5 channel activation. Resting membrane potential was unaffected in *Kcnq5*^−/−^ mutant mice (Fig. 2 a,b,c). Notably, phenylephrine (PE)-induced depolarization of the resting potential was unaffected by *Kcnq5* disruption (Fig. 2 d,e,f). However, lack of KCNQ5 channels attenuated SMC membrane hyperpolarization and muscle relaxation caused by flupirtine (Fig. 2 d,e,f,g,h,i,j.k). Together, these results demonstrate that specifically KCNQ5 plays a role in the anti-contractile effects of PVAT, i.e., in adipose-vascular coupling. Unless systemic physiological control mechanisms can compensate for the increased arterial tone, arterial blood pressure should be elevated in mice lacking KCNQ5 but not KCNQ4 channels. The mean arterial blood pressure of the KCNQ5 knockout mice was indeed elevated. Blood pressure telemetry revealed higher systolic, diastolic, and mean arterial pressure (MAP) in *Kcnq5*^−/−^ and *Kcnq5*^dn/dn^, but not in *Kcnq4*^−/−^ mice (Fig. 1 g,h; Fig. S7). We observed no changes in baroreflex function across all genotypes (Fig. S8).

**Fig. 2.**
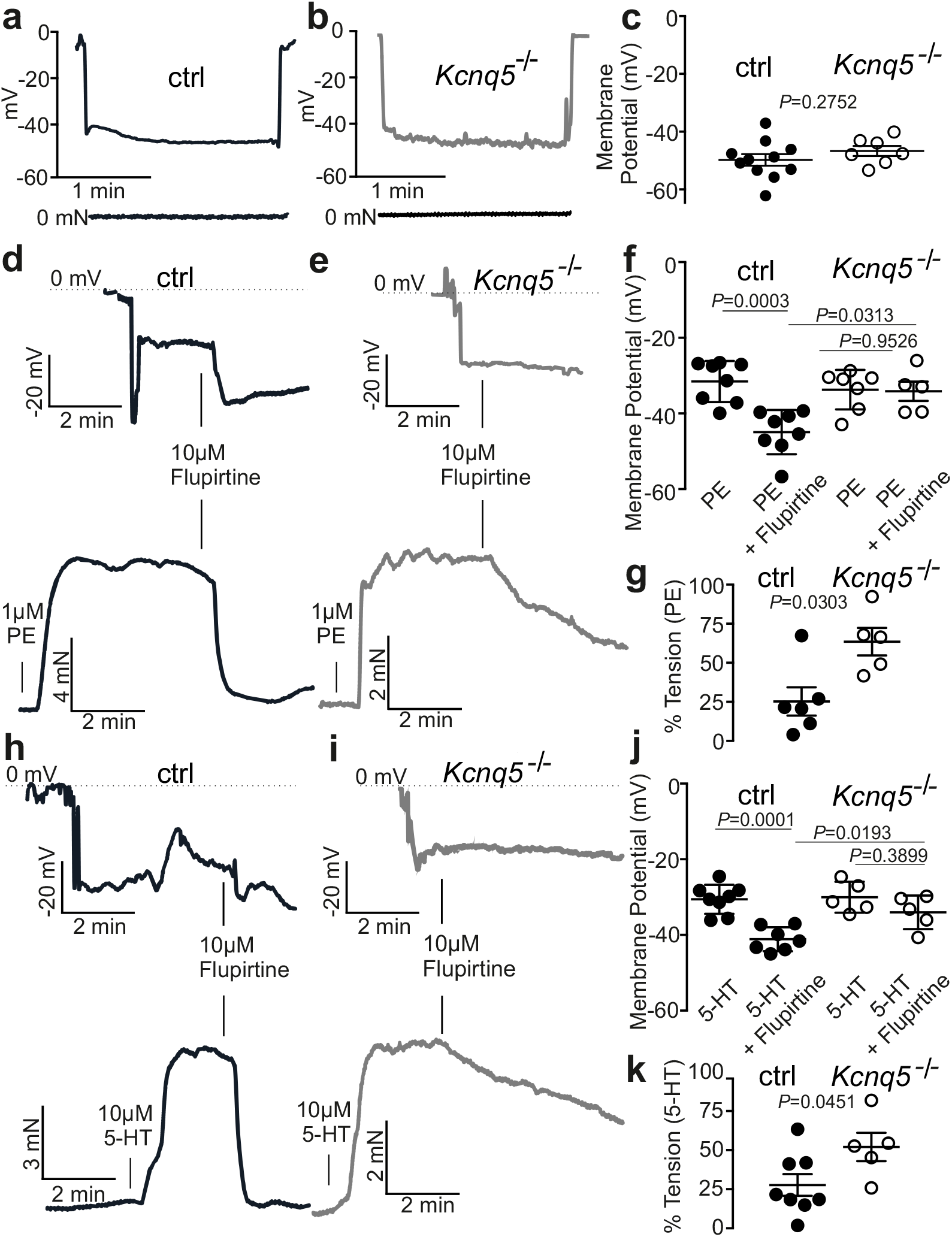
KCNQ5 activation hyperpolarizes smooth muscle cells in intact arteries. Sharp-electrode recordings of the smooth muscle membrane potential in mesenteric arteries isolated for control (*Kcnq5*^+/+^) (**a**) and *Kcnq5*^−/−^ mice (**b**), (**c**). Simultaneous membrane potential and force recordings of vessels treated with alpha 1-adrenoreceptor agonist and subsequent flupirtine. Arteries were isolated from control (*Kcnq5*^+/+^) (**d**) and *Kcnq5*^−/−^ (**e**) mice. Alpha 1-adrenoreceptor agonist induced membrane potential depolarization whereas flupirtine induced membrane potential hyperpolarization in control (*Kcnq5*^+/+^) and *Kcnq5*^−/−^ arteries (**f**). Flupirtine-induced relaxation in control (*Kcnq5*^+/+^) and *Kcnq5*^−/−^ arteries (**g**). Simultaneous membrane potential and force recordings of 5-HT and subsequent flupirtine treated arteries isolated from control (*Kcnq5*^+/+^) (**h**) and *Kcnq5*^−/−^ (**i**) mice. 5-HT-induced depolarization and flupirtine-induced hyperpolarization in control (*Kcnq5*^+/+^) and *Kcnq5*^−/−^ arteries (**j**). Flupirtine-induced relaxation in control (*Kcnq5*^+/+^) and *Kcnq5*^−/−^ arteries (**k**). Data are mean and s.e.m. *P* as determined by ANOVA with Šidák post hoc test.

### Perivascular relaxing factors

Aging is an independent risk factor for hypertension, cardiovascular morbidity, and mortality. We recently demonstrated impaired PVAT control of vascular tone particularly with increased age through metabolic and inflammatory processes and release of perivascular adipose tissue-derived relaxation factors (PVRFs, or ADRF).^32^ Since our results revealed a key effector function of KCNQ5 channels integrating PVAT signals to vasorelaxation, we used targeted lipidomics profiling to characterize the KCNQ5-PVRFs pathway further. This approach identified numerous oxylipins, such as 12,13-DiHOME; 9,10-DiHOME; 9-oxo-ODE; trans-EKODE; 9-HOTrE; 5-HETE; 17,18-DiHETE; 14,15-DHET; 5,6-EET; PGF1a 6-keto; PGD2; and LXB4 as being released into the bath solution from PVAT from young adult mice (10 weeks). Secretion of a number of these oxylipins was reduced with increased age (60 weeks) (Fig. 3a). The mixture of the oxylipins released by PVAT from young mice produced relaxation of arterial rings without PVAT (Fig. 3 b,c,d,f). Importantly, the mixture caused arterial SMC membrane hyperpolarization and relaxation, which were reversed by the pan-KCNQ channel inhibitor XE991 (Fig. 3 d,e,g). Notably, the mixture of oxylipins (e.g. 12,13-DiHOME, 9,10-DiHOME, 9-oxo-ODE) and COX (prostaglandins D2, 6-keto-prostaglandin F2alpha) products released from vessels of aged mice caused transient relaxations and subsequent contractions of young vessels (Fig. 3 b,c). Secretion of the oxylipins is largely independent of alpha-1 adrenoreceptor or 5-HT receptor stimulation (Table S1). We tested the vasodilatory effects of the eight identified oxylipins as mixture and individually. We found that the vasorelaxant effects of the lipid mixture could not be predicted from individual effects of its components^33^; i.e. the individual oxylipins potentiated each other’s vasodilatory effects to cause global relaxation (Fig. S9). In some of the experiments we took advantage of the rat aorta model that was originally used to discover PVAT control of vascular tone ^31 4^ and which remains a valuable model for PVAT studies.^5^ Importantly, the relaxant effects of the mixture were not observed in *Kcnq5*^−/−^ mesenteric arteries (Fig. 3 h), demonstrating an essential role of KCNQ5 channels in the vasodilatory effects of the identified oxylipins (PVRFs). We speculate that the mixture of oxylipins (PVRFs) activates KCNQ5 channels indirectly, namely without direct binding to the KCNQ5 channel protein, and on the anatomical integrity of the vessels. Our opinion is based on the fact that the oxylipin mixture failed to activate KCNQ channels in isolated SMCs and hetero logolouly expressed KCNQ5 channels either in HEK293 cells or in *Xenopus* oocytes, but enabled SMC hyperpolarization in intact tissue. This state of affairs contrasts to effects of high concentration of docosahexaenoic acid (DHA) (Fig. S10), which was shown to directly activate KCNQ5 channels through electrostatic interactions with positively charged gating charges in the upper half of the voltage-sensing domain.^34^

**Fig. 3.**
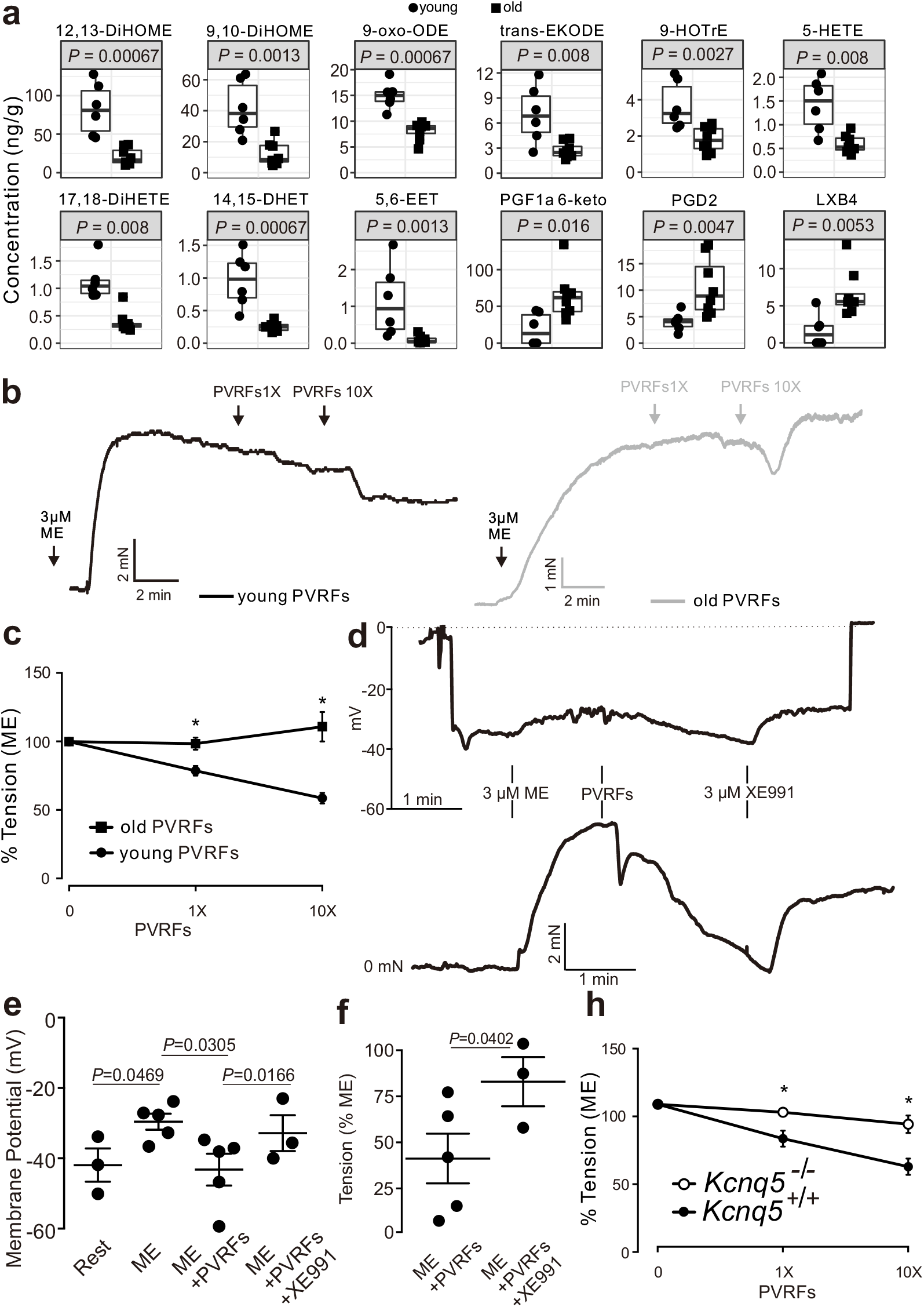
Identification of PVAT-derived relaxation factors. PVAT-derived relaxing factors (PVRFs) released from PVAT of young and old mice **(a)**. N≥6. Relaxation of (−) PVAT mesenteric arteries by synthetic oxylipins in concentrations found after incubation with PVAT of young or old mice **(b, c)**. Molar ratios were used. n≥10 arteries from N=5 mice. Simultaneous membrane potential and force recordings of alpha1 adrenoreceptor agonist, and subsequent PVRFs and XE991 treated arteries **(d)**. Alpha1-adrenoreceptor agonist-induced membrane depolarization, PVRFs-induced membrane hyperpolarization and XE991-induced membrane depolarization of mesenteric arterial smooth muscle cells **(e)**. PVRFs-induced relaxation and methoxamine (ME) and XE991 induced contraction in control arteries **(f)**. PVRFs-induced relaxation in control (*Kcnq5*^+/+^) and *Kcnq5*^−/−^ arteries **(h)**. n≥5 arteries from N≥3 mice.

### Discussion

An influence of PVAT on arterial tone and blood pressure has been hypothesized for decades. We show that PVAT signaling is mediated by KCNQ5 channels, which we now identify as a potential therapeutic blood pressure-lowering target. Here, we used a wide range of *Kcnq* mouse models to indicate a unique role of KCNQ5 in vasoregulation. KCNQ5 is an essential element in adipose-vascular coupling in small resistance vessels and, therefore, in blood pressure regulation. The pathway seems to be specific for coupling perivascular adipose tissue activity to vasodilation, termed the “outside-to-inside” paradigm.^35^ The mechanism was not influenced by cAMP/NO-cGMP signaling pathways, which are involved in endothelial relaxation, i.e. in the “inside-to-outside” control of vascular tone.^35^ Instead, KCNQ5 channels in SMCs facilitate arterial relaxation in response to pharmacological interventions, such as synthetic KCNQ activators, oxylipins, and other PVRFs released by PVAT. Interestingly, hypotensive botanical “folk medicine” has been suggested to regulate KCNQ5 potassium channels in blood vessels.^36^ Our conclusion is further supported by our previous study showing that myogenic contractions persist in arteries deficient in various KCNQ channels, including KCNQ5.^20^

The possible involvement of potassium channels in the control of arterial vascular tone by PVAT has been suggested for decades.^4^ However, convincing data were lacking. Although few studies have reported that Ca^2+^-activated maxi-K^+^ channels (BK_Ca_) might be involved, ^37 38^ other studies have failed to demonstrate their involvement, particularly when genetically modified mouse models were examined.^42^ Intermediate (IK)- and small (SK)-conductance Ca^2+^-sensitive potassium channels are unlikely involved in PVAT control of arterial tone. ^28^ On the basis of pharmacological tools, we previously suggested that voltage-gated KCNQ (K_v7_) channels might be involved.^28 39^ However, because KCNQ blockers and activators have a suspiciously broad spectrum of action in arterial tissue,^40^ possible off-target effects raising doubts about the involvement of KCNQ channels. Using a wide range of genetic mouse models targeting *Kcnq3, Kcnq4* and *Kcnq5* genes (*Kcnq3*^−/−^, *Kcnq4*^−/−^, *Kcnq5*^−/−^, *Kcnq5*^dn/dn^, *Kcnq4*^−/−^*/Kcnq5*^dn/dn^, *Kcnq4*^−/−^*/Kcnq5*^−/−^), we were able to demonstrate a unique role of KCNQ5 in PVAT control of arterial tone. We were able to demonstrate SMC hyperpolarization by synthetic KCNQ channel openers causing relaxation of mesenteric arteries of wild-type but not *Kcnq5*^-/-^ mice. We previously found that PVAT control of arterial tone is normal in *Kcnq1*^-/-^ mice. ^21^ Together, our results provide definitive evidence that KCNQ channel opening in SMCs can lead to relaxation of mouse mesenteric arteries. The signal relies on the KCNQ5 subtype to enact perivascular adipose tissue-mediated vasodilation, and decreased vascular resistance and blood pressure. Although KCNQ2, KCNQ3 and KCNQ5 channels have been proposed provide a hyperpolarizing influence in baroreceptor neurons of rats, ^41^ baroreflex function was normal in in *Kcnq5*^−/−^, *Kcnq5*^dn/dn^ and *Kcnq4*^−/−^ mice.

The identity of PVRFs underling PVAT control of arterial tone is largely unclear. Although few studies have reported that adiponectin might be involved, ^37 38^ other studies have failed to demonstrate its involvement, particularly when genetically modified mouse models were examined.^42^ *Gas-transmitters*, i.e. hydrogen sulfide (H_2_S) and nitric oxide (NO), have been proposed to play a role in adipose-vascular coupling. However, lack of cystathionine γ-lyase did not influence the anticontractile effects of PVAT. ^43^ Furthermore, L-NAME treatment of vessels did not influence the anticontractile effects of PVAT in endothelium-denuded *Cth*^+/+^ arteries.^43^ In this study, we used targeted lipidomics profiling to identify novel candidate PVFRs. This approach identified numerous oxylipins and COX products that together caused relaxation of wild-type, but not of *Kcnq5*^-/-^ mesenteric arteries *via* SMC membrane hyperpolarization. Notably, some of the lipid PVRFs tested, caused no relaxation or had little effects on vascular tone at the single-factor level. Our experimental approach does not exclude the possibility that additional PVRFs are involved in adipose-vascular coupling; however, the results do not inspire confidence. Although examining a single active molecule factor may provide mechanistic insights, understanding the global vasodilatory effect requires considering the multitude of factors impinging on the overall contractile state of the vessel determining vascular tone. At the single-factor level, some substances may have neutral, negative, or positive effects on a number of key responses. However, an increasing number of global change factors would be expected to cause directional changes, which can cause greater total effects of multiple factors.^33^ We used this bidirectional global change approach ^33^ to better understand how vessels react when exposed to number of factors targeting a single effector system, namely the KCNQ5 channel in the blood vessel. Together, we identified numerous oxylipins and COX products as important PVRFs targeting this effector system (KCNQ5), to enable PVAT control of arterial tone and blood pressure. Notably, aging affected the oxylipins profile vasodilatory signaling *via* the KCNQ5 pathway.

Several strategies are available to restore the PVAT function in patients with metabolic syndrome and hypertension. Roux-en-Y gastric bypass, diet-induced weight loss, and increased physical activity all have beneficial effects.^44 45 46^ However, these approaches are invasive and/or face a host of compliance-related issues, demonstrating the need for pharmacological interventions. In addition to drug therapy, for example with glucagon-like peptide-1 (GLP-1) agonists or agents targeting inflammatory interleukins ^47 48 49 50^, addressing oxylipins and KCNQ5 channels may improve outcomes further. For instance, a cytochrome P450-derived epoxide of linoleic acid, termed 12,13-DiHOME, is a recently identified exercise-induced lipokine that increases skeletal muscle fatty acid uptake. As a result, cardiac function and hyperlipidemia could be improved.^51 52^

Hypertension is after all, a condition brought about by an increase in peripheral vascular resistance. We identified novel, hitherto unappreciated elements participating in this process. We suggest that perivascular adipose-tissue coupling to vascular function plays an important role on vasoregulation. We identify KCNQ5 as pivotal to that process, which requires the anatomical integrity of the vessels. Our results suggest potential novel targets for the therapeutic interventions focused on PVAT–control of vascular tone in health, cardiovascular disease, and aging.

## Acknowledgements

We thank Ilona Kramer for excellent assistance in telemetric blood pressure measurements, Inci Dogan for help with GC/MS analysis. Wolf-Hagen Schunck supported our studies and gave valued advice. We thank Ning Wang for help in studies on isolated aortas. The Deutsche Forschungsgemeinschaft (DFG) (GO766/15-2, supported our study (GO766/22-3, GO 766/12-3 and SFB 1365) (M.G., T.J.J.) and under Excellence Strategy – EXC-2049-390688087 (NeuroCure) (T.J.J.).

## Data availability

The data that support the findings of this study are available from the corresponding author upon reasonable request.

## Methods

### Mouse Model

We used our previously generated *Kcnq3*^−/−^ mice.^53^ Briefly, exon 3 was removed from the *Kcnq3* locus, which led to a frameshift. In *Kcnq4*^−/−^ mice, exons 6–8 were deleted (leading to a frameshift) described in detail.^17^ *Kcnq4*^−/−^ mice were backcrossed to a C3H background (>10 generations).^54^ The dominant negative G278S point mutation was introduced in *Kcnq5*^dn/dn^ mice as previously reported.^18^ *Kcnq5*^−/−^ mice were generated by introducing 2 stop codons in Exon 6 (TAA TGA).^55^ *Kcnq4*^−/−^/*Kcnq5*^dn/dn^ and *Kcnq4*^−/−^/*Kcnq5*^−/−^ double mutant mice were generated by breeding *Kcnq4*^−/+^/*Kcnq5*^dn/+^ and *Kcnq4*^−/+^/*Kcnq5*^−/+^ mice, respectively. *Kcnq4*^+/−^ /*Kcnq5*^+/–^ double mutant mice were backcrossed to a Bl6 background (>10 generations). Heterozygous mice were used for breeding to obtain homozygous knockout mice. Littermate (age 8–12 weeks) male wild-type mice (^+/+^) were used as controls.

We also used young (age 11–18 weeks), old (age 50–69 weeks) male wild-type mice C57BL/6N. Animal care followed American Physiological Society guidelines, and local authorities (Landesamt für Gesundheit und Soziales Berlin, LAGeSo) approved all protocols. Mice were housed in individually ventilated cages under standardized conditions with an artificial 12-h dark–light cycle with free access to water and food. Animals were randomly assigned to the experimental procedures in accordance with the German legislation on protection of animals.

### Wire Myography

Mesenteric arteries were isolated after sacrifice with isoflurane anesthesia, as previously described.^21 39^ Blood vessels were quickly transferred to cold (4°C), oxygenated (95% O_2_/5% CO_2_) physiological salt solution (PSS) containing (in mmol/L) 119 NaCl, 4.7 KCl, 1.2 KH_2_PO_4_, 25 NaHCO_3_, 1.2 Mg_2_SO_4_, 11.1 glucose, and 1.6 CaCl_2_. We dissected the vessels into 2 mm rings whereby perivascular fat and connective tissue were either intact [(+) PVAT] or removed [(−) PVAT rings]. Each ring was placed between two stainless steel wires (diameter 0.0394mm) in a 5-ml organ bath of a Mulvany Small Vessel Myograph (DMT 610M; Danish Myo Technology, Denmark). The organ bath was filled with PSS. Continuously oxygenated bath solution with a gas mixture of 95% O_2_ and 5% CO_2_ was kept at 37°C (pH 7.4). To obtain the passive diameter of the vessel at 100mm Hg, a DMT normalization procedure was used. The mesenteric artery rings were placed under a tension equivalent to that generated at 0.9 times the diameter of the vessel at 100mm Hg by stepwise distending the vessel using the LabChart DMT Normalization module. The software Chart5 (AD Instruments Ltd. Spechbach, Germany) was used for data acquisition and display. After 60-min incubation, arteries were precontracted either with isotonic external 60 mM KCl or alpha1-adrenoreceptor (methoxamine (ME) or phenylephrine (PE)), serotonin (5–HT), thromboxane A 2 receptor (U46619) agonists, until a stable resting tension was acquired. Alpha-1 adrenoreceptor and 5-HT contractions were similar across all mouse genotypes used in the study (Fig S11). The composition of 60 mM KCl (in mmol/L) was 63.7 NaCl, 60 KCl, 1.2 KH_2_PO_4_, 25 NaHCO_3_, 1.2 Mg_2_SO_4_, 11.1 glucose, and 1.6 CaCl_2_. Drugs were added to the bath solution if not indicated otherwise. Tension is expressed as a percentage of the steady-state tension (100%) obtained with isotonic external 60 mM KCl or an agonist.

### Isometric contractions of rat vessels

Male Sprague-Dawley rats (200-300 g, 8-12 weeks; Charles River, Sulzfeld/Berlin Germany) were killed, and the thoracic aortas were removed, and quickly transferred to cold (4°C) oxygenated (95% O_2_/5% CO_2_) physiological salt solution (PSS), and dissected into 2 mm rings, respectively. Perivascular fat and connective tissue were removed as previously described. ^56^

^57^ After 1h equilibration, contractile force was measured isometrically using standard procedures and solutions as described.^58^ The bath solution volume was 20 mL. Cumulative concentration response curves were obtained for 5-HT agonist (serotonin) in the presence and absence of XE991 inhibitor (10,10-bis(4-pyridinylmethyl)-9(10H)-anthracenone dihydrochloride. Tension was expressed as a percentage of the steady-state tension (100%) obtained with 5-HT agonist.

### Membrane Potential Recordings

Intracellular recordings of membrane potential in smooth muscle cells of intact mesenteric arteries were performed using sharp microelectrodes. They were pulled from aluminosilicate glass using P-97 Flaming/Brown Micropipette Puller (Sutter Instrument, USA). Sharp electrodes were filled with 3M KCl as previously described. ^59 32^ An amplifier (DUO 773, World Precision Instruments) was used to record the membrane potential. We used a micromanipulator (UMP, Sensapex, Finnland) to make impalements from the vessel’s adventitial side. The following criteria for acceptance of membrane potential recordings were used: (1) an abrupt change in membrane potential upon cell penetration; (2) a constant electrode resistance when compared before, during, and after the measurement; (3) a stable reading of the membrane potential lasting longer than 1 min; and (4) no change in the baseline when the electrode was removed.

### Two-Electrode Voltage Clamp Recordings

KCNQ5 cDNA was inserted into pTPN vector (internal database number 1270)^60^ and myc-tagged KCNQ4 cDNA was inserted into pTLN (1491). Vectors were linearized with HpaI, and capped cRNA was transcribed using SP6 mMessage mMachine kit (Ambion). *Xenopus* oocytes were defolliculated by collagenase treatment, injected with 13 ng (KCNQ5) or 28 ng (KCNQ4) of cRNA and incubated for 2 days at 17 °C in penicillin/streptomycin-containing saline. Measurements were performed at room temperature in ND109 solution (109 mM NaCl, 2 mM KCl, 1.8 mM CaCl_2_, 1 mM MgCl_2_, 5 mM HEPES-NaOH, pH 7.4) using a Turbotec (npi Instruments) amplifier and pClamp 10.7 (Axon Instruments) software. Voltage was clamped at −80 mV, and stepped every 5 s to voltages between −100 and +60 mV for 2 s in 10 mV increments followed by a 500-ms 30 mV step. Current was measured at the end of each 2-s voltage step.

### Patch-Clamp Recordings

Potassium currents were recorded in the conventional whole-cell configuration of the patch-clamp technique at room temperature using an Axopatch 200B (Axon Instruments/Molecular Devices, Sunnyvale, CA) or an EPC7 (List, Darmstadt, Germany) amplifier under control of Clampex software (Molecular Devices, Sunnyvale, CA) and digitized at 5 kHz using a Digidata 1440A digitizer (Axon CNS, Molecular Devices). ^61 62^ Patch pipettes were made from borosilicate glass capillary tubes, had resistances of 2 to 5 MΩ, and were filled with solution containing (in mmol/L) 130 KCl, 1 MgCl_2_, 3 Na_2_ATP, 0.1 Na_3_GTP, 10 HEPES, and 5 EGTA (pH, 7.2). The external solution contained (in mmol/L) 126 NaCl, 5 KCl, 1 MgCl_2_, 0.1 CaCl_2_, 11 glucose, and 10 HEPES (pH, 7.2). Time courses of outward currents at +20 mV from a holding potential of −60 mV were recorded from transfected HEK293 cells during extracellular application of PVRFs or DHA (70 µM).

### Radiotelemetric blood pressure measurement

Mice were anesthetized by intraperitoneal application of ketamine/xylazine adapted to body weight. Miniature subcutaneous radiotelemetry transmitters (PhysioTel PA-C10, Data Sciences International New Brighton, MN, USA) were implanted into the left carotid artery as previously described. ^63 64 3^ After 10 days of recovery from surgery, interventions and/or recordings (Dataquest A.R.T. software for acquisition and analysis) were started. Systolic, diastolic blood pressures, and heart rate were recorded continuously in freely moving mice. Nocturnal blood pressure patterns were normal (data are not shown). To test baroreflex, 2 mg atropine, 4 mg L-NAME, 4 and 8 mg metoprolol, 1 mg prazosin, were administered intraperitonially (i.p). All drugs were solved in sterile PSS. Blood pressures were recorded during 1 hour at baseline and 1 hour after drugs administration between 9 AM and 11 AM. Atropine for muscarinic blockade and prazosin for alpha1-adrenergic receptor blockade were used to study the role of KCNQ channels in the autonomic BP control ^43^.

### RNA Sequencing

Following Agilent 2100 bioanalyzer quality control, RNA-seq was performed using Illumina Genome Analyzer Novaseq 6000 platform. NEB Next® Ultra™ RNA Library Prep Kit was used for library preparation. Sequence quality estimations, GC content, nucleotide distribution, and read duplication levels were determined for the samples using fastp-0.12.2 software. The reads were mapped to the reference mouse genome (ensembl_mus_musculus_grcm38_p6_gca_000001635_8). HISAT2 was selected to map the filtered sequenced reads to the reference genome. The uniquely mapped read data output was processed using custom scripts in R software (version 3.5.1) and then normalized using the FeatureCounts package v1.5.0-p3 version. Differential expression analysis was performed using the DESeq2 R package version v1.20.0.^65^ We used clusterProfiler for enrichment analysis, including GO Enrichment, DO Enrichment, Kyoto Encyclopedia of Genes and Genomes (KEGG), and Reactome database Enrichment.^66^

## Materials

All salts and other chemicals were purchased from Sigma-Aldrich (Germany) or Merck (Germany). Using DMSO or PSS, drugs were freshly dissolved on the day of each experiment accordingly to the material sheet. Maximal DMSO concentration after application did not exceed 0.5%.

## Statistics

Data are presented as mean±SEM. We calculated EC50 values using a Hill equation: T=(B0 – Be)/(1+([D]/EC50)n)+Be, where T is the tension in response to the drug (D); Be is the maximum response induced by the drug; B0, is a constant; EC50 is the concentration of the drug that elicits a half-maximal response. For curve fittings using non-linear regression, GraphPad 8.0.1 (Graphpad Software, San Diego, CA, United States) software was used. Statistical significance was determined by Mann–Whitney test or nonparametric ANOVA (Kruskal-Wallis test). Extra sum-of-squares F test was performed for comparison of concentration-response curves. Values of p<0.05 were considered statistically significant. n represents the number of arteries; N represents the number of mice tested. We used at least n=5 from at least N=3 for wire myograph experiments. Based on experimental data, we determined the sample size and power of statistical analyses to be optimal for such experiments. ^67^ Figures were made using CorelDraw Graphics Suite 2020 (Corel Corporation Ottawa, Canada).

## Lipidomics profiling

### Gas chromatography/mass spectrometry (GC/MS) analysis

Lipid measurements were performed by Lipidomics (Berlin, Germany). GC/MS analysis was performed using an Agilent ChemStation. ^68 69 70^ For determination of PVRFs concentrations, rat peri-aortic adipose tissue (1 g) was incubated in 15 mL-Eppendorf tubes with 10 mL PSS solutions. After removal of adipose tissue, the PSS solution were dissolved in hexane, extracted and vortexed. Next, 1 ml water was added to the solution. In order to ensure that the concentration between the aqueous and the lipophilic phase was in equilibrium the samples were shaken by hand for 4 min. The phases were then separated by centrifugation and the lipophilic hexane phase containing fatty acid methyl esters was removed and dried under nitrogen. The fatty acid methyl ester residues were redissolved in 50 µL hexane and transferred into an autosampler vial. Samples were analyzed by using a fully automated Agilent 7890A-5977B system equipped with a flame ionization detector. Peaks of resolved substances were identified by comparison with a standard.

### Sample preparation for analysis of PUFA-derived lipid mediators in mice

Briefly, 500 µL bath solution were mixed with internal standard compounds (14,15-DHET-D11; 15-HETE-d8; 20-HETE-d6; 8,9-EET-d11; 9,10-DiHOME-d4; d4-12(13)-EpOME; d4-13-HODE; d4-PGB2; d4-PGE2-13,14-dihydro-15-keto; d4-PGF2a; LTB4-D4; PGE2-D4; from Cayman Chemicals, Ann Arbor, MI, USA) and directly extracted with methanol/water followed by solid-phase extraction of the metabolites. Methanol was added for protein precipitation. After centrifugation and pH adjustment at 6.0, the obtained supernatant was added to Bond Elute Certify II columns (Agilent Technologies, Santa Clara, USA) for Solid phase Extraction. The eluate was evaporated on a heating block at 40 °C under a stream of nitrogen to obtain a solid residues that were dissolved in 100 µL methanol/water.

### LC/ESI-MS/MS

The residues were analyzed using an Agilent 1290 HPLC system with binary pump, multisampler and column thermostat with a Zorbax Eclipse plus C-18, 2.1 × 150 mm, 1.8 µm column using a gradient solvent system of aqueous acetic acid (0.05%) and acetonitrile / methanol 50:50. The flow rate was set at 0.3 mL/min, the injection volume was 20 µL. The HPLC was coupled with an Agilent 6495 Triplequad mass spectrometer (Agilent Technologies, Santa Clara, USA) with electrospray ionisation source. Analysis was performed with Multiple Reaction Monitoring in negative mode, at least two mass transitions for each compound.^68 69^ _70_

