## Supplemental Material for "KCNQ5 controls perivascular adipose tissue-mediated vasodilation"

Running head: KCNQ5

^1^Dmitry Tsvetkov, ^2^Johanna Schleifenbaum, ^3^Yibin Wang, ^1^Mario Kassmann, ^4,5^Maya M Polovitskaya, ^4,5^Sebastian Schütze, ^6^Michael Rothe,

^3^Friedrich C Luft, ^4,5,7^Thomas J Jentsch, ^1,3^Maik Gollasch

1. University Medicine Greifswald, Department of Internal Medicine and Geriatrics, Ferdinand Sauerbruch Street, Greifswald, Germany
2. Institute of Vegetative Physiology, Charité-Universitätsmedizin Berlin, Berlin, Germany
3. Experimental and Clinical Research Center (ECRC), a joint cooperation between the Charité Medical Faculty and the Max Delbrück Center for Molecular Medicine (MDC), Berlin, Germany
4. Leibniz-Forschungsinstitut für Molekulare Pharmakologie (FMP), Berlin, Germany
5. Max Delbrück Center for Molecular Medicine (MDC) in the Helmholtz Association, Berlin, Germany
6. LIPIDOMIX GmbH, Berlin, Germany
7. NeuroCure Cluster of Excellence, Charité Universitätsmedizin, Berlin, Germany

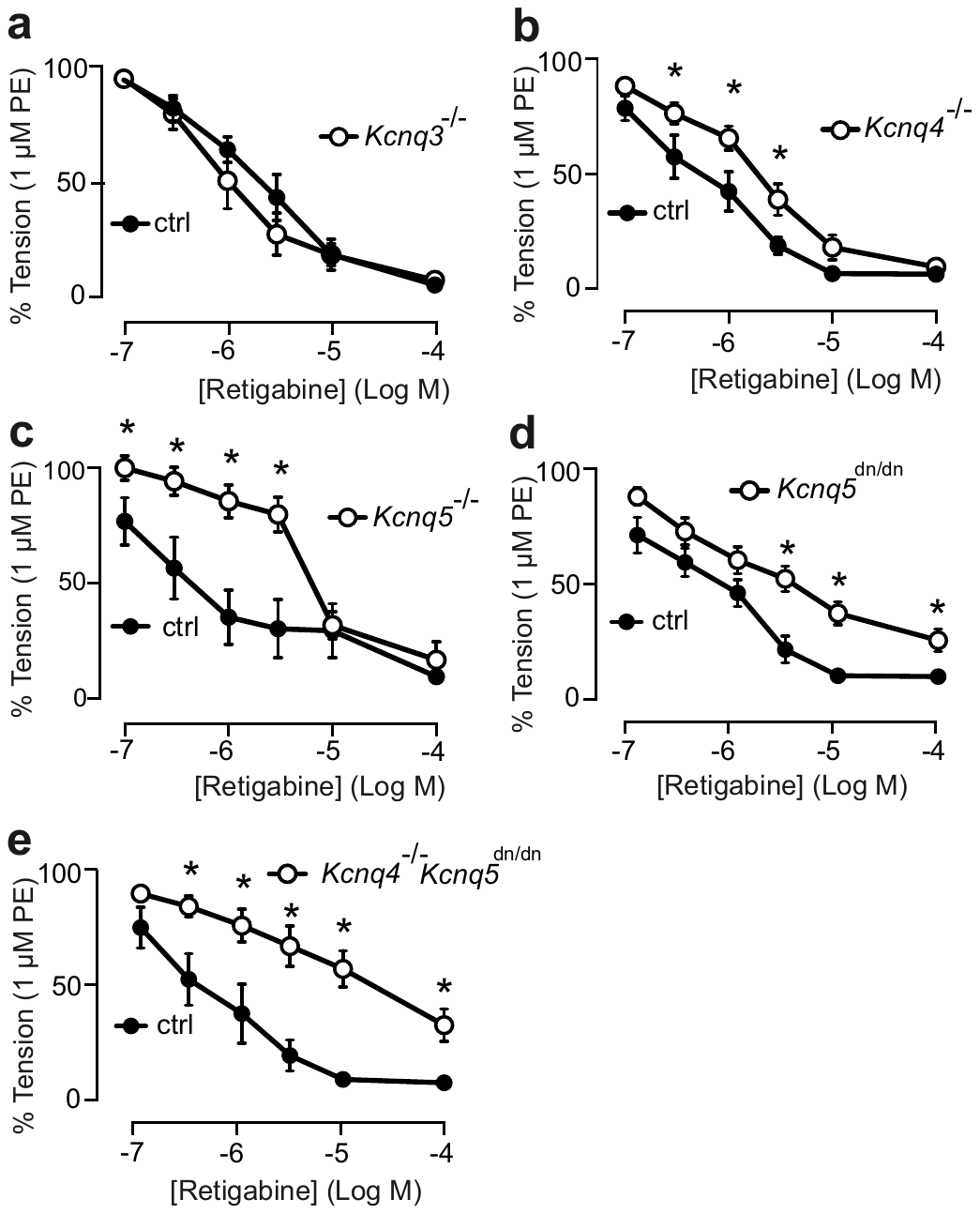

Fig. 1 **KCNQ4 and KCNQ5 but not KCNQ3 channels regulate arterial tone in alpha1-adrenergic agonist precontracted vessels**

ation of () fat mesenteric artery rings by 0.3-3 μM R-L3, 0.1-10 μM ML 277 or 0.01-10 μM re tigabine. Vessels were isolated from wild-t ype

(Kcnq1

+/+

)mice(A)orKcnq1

/

mice (B). Mesenteric arteries were precontracted by phenylephrine (PE) 1 μM (A,B) or KCl 60 mM (A). Represen-

tative traces showing relaxation induced by retigabine in rings from Kcnq1

+/+

(E) and Kcnq1

/

mice (F) and average values compared with vehicle

(DMSO) and water controls (C). ML277 effects on rings from Kcnq1

/

and Kcnq1

+/+

mice (D). Tension is expressed as a percentage of KCl or

PE-induced contractions. n =5pergroup

ation of () fat mesenteric artery rings by 0.3-3 μM R-L3, 0.1-10 μM ML 277 or 0.01-10 μM re tigabine. Vessels were isolated from wild-t ype

(Kcnq1

+/+

)mice(A)orKcnq1

/

mice (B). Mesenteric arteries were precontracted by phenylephrine (PE) 1 μM (A,B) or KCl 60 mM (A). Represen-

tative traces showing relaxation induced by retigabine in rings from Kcnq1

+/+

(E) and Kcnq1

/

mice (F) and average values compared with vehicle

(DMSO) and water controls (C). ML277 effects on rings from Kcnq1

/

and Kcnq1

+/+

mice (D). Tension is expressed as a percentage of KCl or

PE-induced contractions. n =5pergroup

Relaxation of () fat mesenteric artery rings by 0.3-3 μM R-L3, 0.1-10 μM ML 277 or 0.01-10 μM re tigabine. Vessels were isolated from wild-t ype

(Kcnq1

+/+

)mice(A)orKcnq1

/

mice (B). Mesenteric arteries were precontracted by phenylephrine (PE) 1 μM (A,B) or KCl 60 mM (A). Represen-

tative traces showing relaxation induced by retigabine in rings from Kcnq1

+/+

(E) and Kcnq1

/

mice (F) and average values compared with vehicle

(DMSO) and water controls (C). ML277 effects on rings from Kcnq1

/

and Kcnq1

+/+

mice (D). Tension is expressed as a percentage of KCl or

PE-induced contractions. n =5pergrou

Relaxation of () fat mesenteric artery rings by 0.3-3 μM R-L3, 0.1-10 μM ML 277 or 0.01-10 μM re tigabine. Vessels were isolated from wild-t ype

(Kcnq1

+/+

)mice(A)orKcnq1

/

mice (B). Mesenteric arteries were precontracted by phenylephrine (PE) 1 μM (A,B) or KCl 60 mM (A). Represen-

tative traces showing relaxation induced by retigabine in rings from Kcnq1

+/+

(E) and Kcnq1

/

mice (F) and average values compared with vehicle

(DMSO) and water controls (C). ML277 effects on rings from Kcnq1

/

and Kcnq1

+/+

mice (D). Tension is expressed as a percentage of KCl or

PE-induced contractions. n =5pergrou

Relaxation of (−) PVAT mesenteric arteries by retigabine. Vessels were isolated from *Kcnq3*^−/−^ (**a**), *Kcnq4*^−/−^ (**b**) *Kcnq5*^−/−^ (**c**) mice, *Kcnq5*^dn/dn^ (**d**), *Kcnq4*^−/−^*/Kcnq5*^dn/dn^ (**e**), and respective control (*Kcnq3*^+/+^, *Kcnq4*^+/+^, *Kcnq5*^+/+^, *Kcnq4*^+/+^*/Kcnq5*^+/+^) mice. Tension is expressed as a percentage of alpha1 adrenoreceptor agonists-induced contraction. Data are mean ± s.e.m. *P as determined by two-way ANOVA with Šidák post hoc test. n≥5 arteries from N≥3 mice.

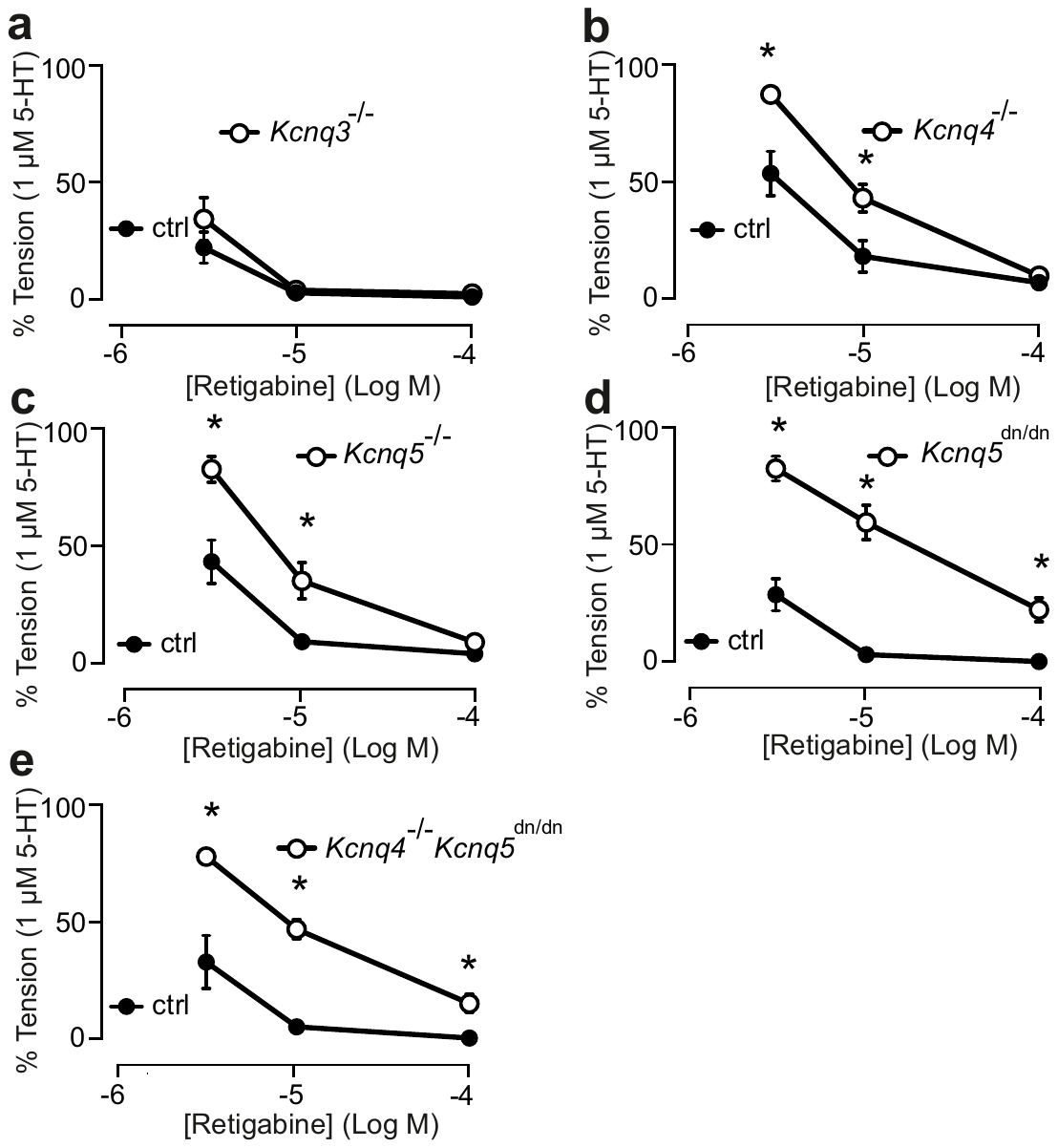

Fig. 2 **KCNQ4/KCNQ5 but not KCNQ3 channels regulate arterial tone in 5-HT precontracted vessels**

Relaxation of (−) PVAT mesenteric arteries by retigabine. Vessels were isolated from *Kcnq3*^−/−^ (**a**), *Kcnq4*^−/−^ (**b**), *Kcnq5*^−/−^ (**c**), *Kcnq5*^dn/dn^ (**d**), *Kcnq4*^−/−^*Kcnq5*^dn/dn^ (**e**) or respective control (*Kcnq3*^+/+^, *Kcnq4*^+/+^, *Kcnq5*^+/+^, *Kcnq4*^+/+^*/Kcnq5*^+/+^) mice. Tension is expressed as a percentage of or 5-HT-induced contraction. 5-HT, serotonin. Data are mean ± s.e.m. *P as determined by two-way ANOVA with Šidák post hoc test. n≥6 arteries from N≥3 mice.

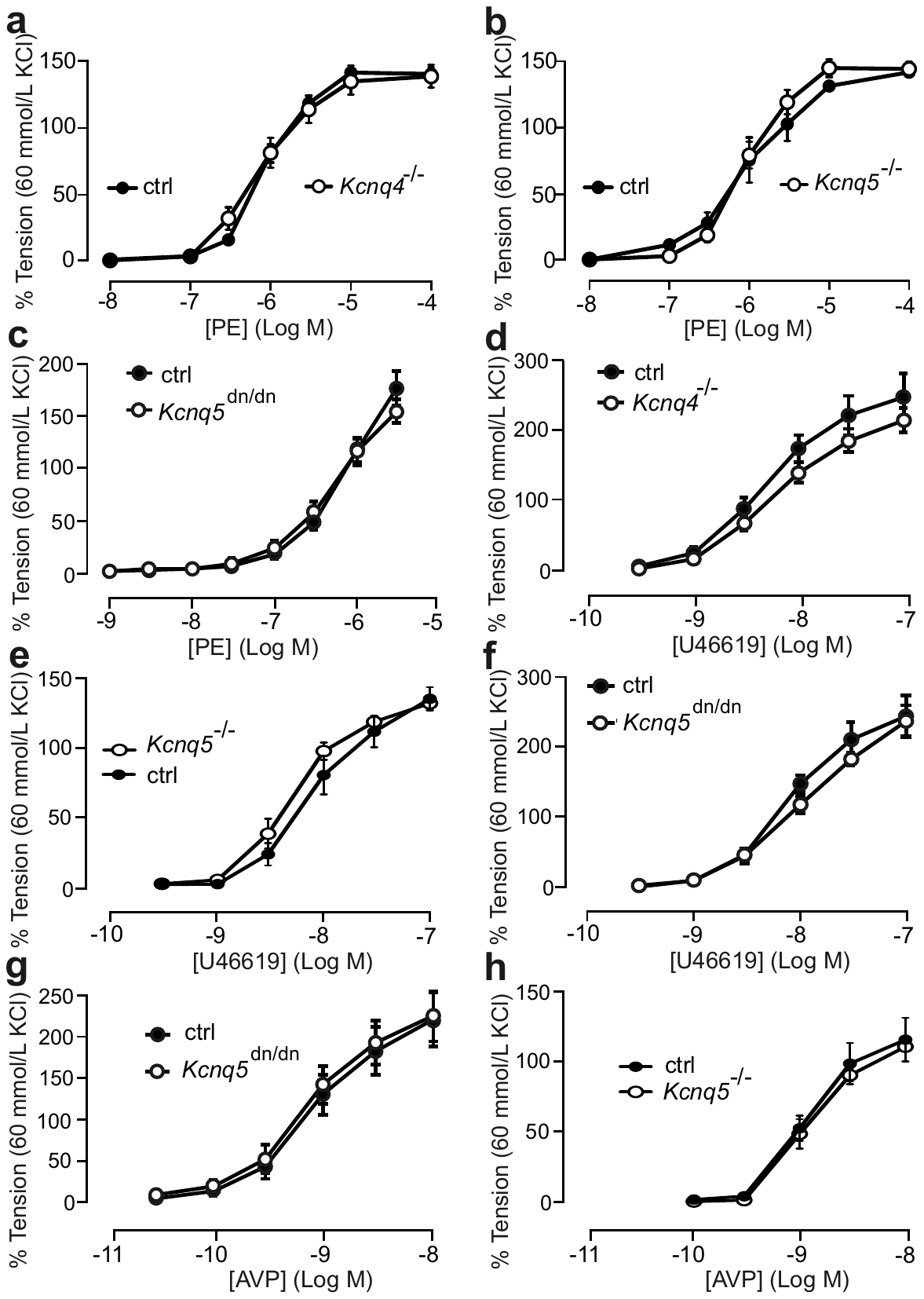

Fig. 3 **KCNQ4 and KCNQ5 channels have no effects in alpha-1 adrenoreceptor, thromboxane A2 receptor agonist, and AVP induced contraction**

Contraction of (−) PVAT mesenteric arteries by a1-adrenoreceptor-agonist in *Kcnq4*^−/−^ (**a**) *Kcnq5*^−/−^ (**b**), *Kcnq5*^dn/dn^ (**c**); thromboxane A2 agonist (U46619) in *Kcnq4*^−/−^ (**d**), *Kcnq5*^−/−^ (**e**), *Kcnq5*^dn/dn^ (**f**); AVP in *Kcnq5*^dn/dn^ (**e**), *Kcnq5*^−/−^ (**f**) mice. Respective *Kcnq4*^+/+^, *Kcnq5*^+/+^ were used as control. AVP, vasopressin. Data are mean ± s.e.m. *P as determined by two-way ANOVA with Šidák post hoc test. n≥6 arteries from N≥3 mice.

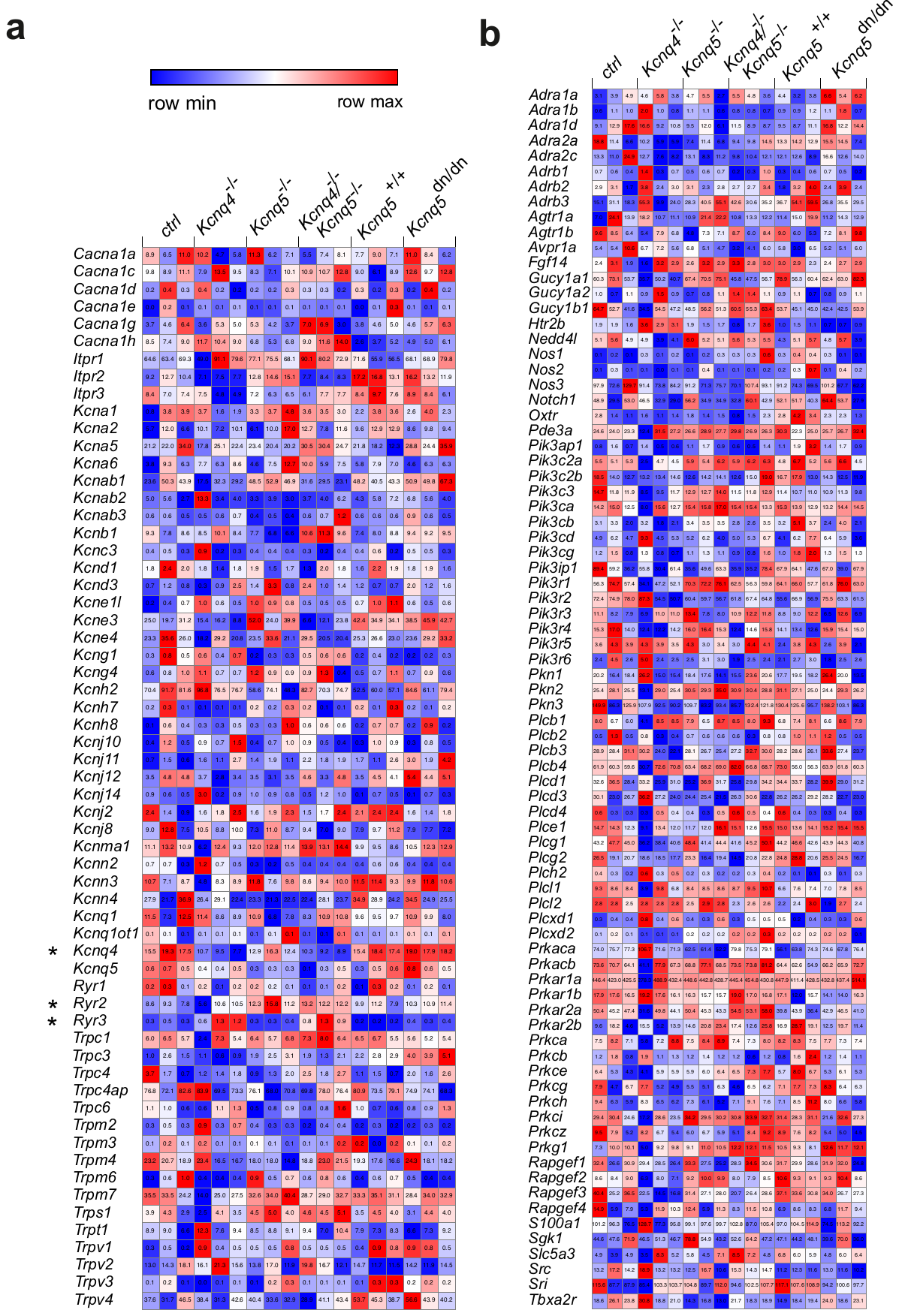

Fig. 4 **RNA-Sequencing did not reveal dysregulation of pathways involved in vascular tone**

Heatmap displaying Fragments Per Kilobase of transcript per Million mapped reads (FPKM) values for mRNA of ion channels (**a**), receptors and second messengers (**b**) in (–) PVAT mesenteric arteries isolated from control, *Kcnq4*^−/−^, *Kcnq5*^−/−^, *Kcnq4*^−/−^/*Kcnq5*^−/−^ mice. Wald test was used. N=3 mice. *adjusted P value < 0.05. *Kcnq4*^−/−^, *Kcnq4*^−/−^/*Kcnq5*^−/−^ mice expressed lower *Kcnq4*. *Ryr3* expression was higher in *Kcnq4*^−/−^ and *Kcnq4*^−/−^/*Kcnq5*^−/−^ mice vs. control. *Ryr2* was overexpressed only in *Kcnq4*^−/−^/*Kcnq5*^−/−^. *Kcnq5*^dn/dn^ did not show such findings.

| Gene Name | Gene Description |
| --- | --- |
| *Cacna1a* | calcium channel, voltage-dependent, P/Q type, alpha 1A subunit [Source:MGI Symbol;Acc:MGI:109482] |
| *Cacna1c* | calcium channel, voltage-dependent, L type, alpha 1C subunit [Source:MGI Symbol;Acc:MGI:103013] |
| *Cacna1d* | calcium channel, voltage-dependent, L type, alpha 1D subunit [Source:MGI Symbol;Acc:MGI:88293] |
| *Cacna1e* | calcium channel, voltage-dependent, R type, alpha 1E subunit [Source:MGI Symbol;Acc:MGI:106217] |
| *Cacna1g* | calcium channel, voltage-dependent, T type, alpha 1G subunit [Source:MGI Symbol;Acc:MGI:1201678] |
| *Cacna1h* | calcium channel, voltage-dependent, T type, alpha 1H subunit [Source:MGI Symbol;Acc:MGI:1928842] |
| *Itpr1* | inositol 1,4,5-trisphosphate receptor 1 [Source:MGI Symbol;Acc:MGI:96623] |
| *Itpr2* | inositol 1,4,5-triphosphate receptor 2 [Source:MGI Symbol;Acc:MGI:99418] |
| *Itpr3* | inositol 1,4,5-triphosphate receptor 3 [Source:MGI Symbol;Acc:MGI:96624] |
| *Kcna1* | potassium voltage-gated channel, shaker-related subfamily, member 1 [Source:MGI Symbol;Acc:MGI:96654] |
| *Kcna2* | potassium voltage-gated channel, shaker-related subfamily, member 2 [Source:MGI Symbol;Acc:MGI:96659] |
| *Kcna5* | potassium voltage-gated channel, shaker-related subfamily, member 5 [Source:MGI Symbol;Acc:MGI:96662] |
| *Kcna6* | potassium voltage-gated channel, shaker-related, subfamily, member 6 [Source:MGI Symbol;Acc:MGI:96663] |
| *Kcnab1* | potassium voltage-gated channel, shaker-related subfamily, beta member 1 [Source:MGI Symbol;Acc:MGI:109155] |
| *Kcnab2* | potassium voltage-gated channel, shaker-related subfamily, beta member 2 [Source:MGI Symbol;Acc:MGI:109239] |
| *Kcnab3* | potassium voltage-gated channel, shaker-related subfamily, beta member 3 [Source:MGI Symbol;Acc:MGI:1336208] |
| *Kcnb1* | potassium voltage gated channel, Shab-related subfamily, member 1 [Source:MGI Symbol;Acc:MGI:96666] |
| *Kcnc3* | potassium voltage gated channel, Shaw-related subfamily, member 3 [Source:MGI Symbol;Acc:MGI:96669] |
| *Kcnd1* | potassium voltage-gated channel, Shal-related family, member 1 [Source:MGI Symbol;Acc:MGI:96671] |
| *Kcnd3* | potassium voltage-gated channel, Shal-related family, member 3 [Source:MGI Symbol;Acc:MGI:1928743] |
| *Kcne1l* | potassium voltage-gated channel, Isk-related family, member 1-like, pseudogene [Source:MGI Symbol;Acc:MGI:1913490] |
| *Kcne3* | potassium voltage-gated channel, Isk-related subfamily, gene 3 [Source:MGI Symbol;Acc:MGI:1891124] |
| *Kcne4* | potassium voltage-gated channel, Isk-related subfamily, gene 4 [Source:MGI Symbol;Acc:MGI:1891125] |
| *Kcng1* | potassium voltage-gated channel, subfamily G, member 1 [Source:MGI Symbol;Acc:MGI:3616086] |
| *Kcng4* | potassium voltage-gated channel, subfamily G, member 4 [Source:MGI Symbol;Acc:MGI:1913983] |
| *Kcnh2* | potassium voltage-gated channel, subfamily H (eag-related), member 2 [Source:MGI Symbol;Acc:MGI:1341722] |
| *Kcnh7* | potassium voltage-gated channel, subfamily H (eag-related), member 7 [Source:MGI Symbol;Acc:MGI:2159566] |
| *Kcnh8* | potassium voltage-gated channel, subfamily H (eag-related), member 8 [Source:MGI Symbol;Acc:MGI:2445160] |
| *Kcnj10* | potassium inwardly-rectifying channel, subfamily J, member 10 [Source:MGI Symbol;Acc:MGI:1194504] |
| *Kcnj11* | potassium inwardly rectifying channel, subfamily J, member 11 [Source:MGI Symbol;Acc:MGI:107501] |
| *Kcnj12* | potassium inwardly-rectifying channel, subfamily J, member 12 [Source:MGI Symbol;Acc:MGI:108495] |
| *Kcnj14* | potassium inwardly-rectifying channel, subfamily J, member 14 [Source:MGI Symbol;Acc:MGI:2384820] |
| *Kcnj2* | potassium inwardly-rectifying channel, subfamily J, member 2 [Source:MGI Symbol;Acc:MGI:104744] |
| *Kcnj8* | potassium inwardly-rectifying channel, subfamily J, member 8 [Source:MGI Symbol;Acc:MGI:1100508] |
| *Kcnma1* | potassium large conductance calcium-activated channel, subfamily M, alpha member 1 [Source:MGI Symbol;Acc:MGI:99923] |
| *Kcnn2* | potassium intermediate/small conductance calcium-activated channel, subfamily N, member 2 [Source:MGI Symbol;Acc:MGI:2153182] |
| *Kcnn3* | potassium intermediate/small conductance calcium-activated channel, subfamily N, member 3 [Source:MGI Symbol;Acc:MGI:2153183] |
| *Kcnn4* | potassium intermediate/small conductance calcium-activated channel, subfamily N, member 4 [Source:MGI Symbol;Acc:MGI:1277957] |
| *Kcnq1* | potassium voltage-gated channel, subfamily Q, member 1 [Source:MGI Symbol;Acc:MGI:108083] |
| *Kcnq1ot1* | KCNQ1 overlapping transcript 1 [Source:MGI Symbol;Acc:MGI:1926855] |
| *Kcnq4* | potassium voltage-gated channel, subfamily Q, member 4 [Source:MGI Symbol;Acc:MGI:1926803] |
| *Kcnq5* | potassium voltage-gated channel, subfamily Q, member 5 [Source:MGI Symbol;Acc:MGI:1924937] |
| *Ryr1* | ryanodine receptor 1, skeletal muscle [Source:MGI Symbol;Acc:MGI:99659] |
| *Ryr2* | ryanodine receptor 2, cardiac [Source:MGI Symbol;Acc:MGI:99685] |
| *Ryr3* | ryanodine receptor 3 [Source:MGI Symbol;Acc:MGI:99684] |
| *Trpc1* | transient receptor potential cation channel, subfamily C, member 1 [Source:MGI Symbol;Acc:MGI:109528] |
| *Trpc3* | transient receptor potential cation channel, subfamily C, member 3 [Source:MGI Symbol;Acc:MGI:109526] |
| *Trpc4* | transient receptor potential cation channel, subfamily C, member 4 [Source:MGI Symbol;Acc:MGI:109525] |
| *Trpc4ap* | transient receptor potential cation channel, subfamily C, member 4 associated protein [Source:MGI Symbol;Acc:MGI:1930751] |
| *Trpc6* | transient receptor potential cation channel, subfamily C, member 6 [Source:MGI Symbol;Acc:MGI:109523] |
| *Trpm2* | transient receptor potential cation channel, subfamily M, member 2 [Source:MGI Symbol;Acc:MGI:1351901] |
| *Trpm3* | transient receptor potential cation channel, subfamily M, member 3 [Source:MGI Symbol;Acc:MGI:2443101] |
| *Trpm4* | transient receptor potential cation channel, subfamily M, member 4 [Source:MGI Symbol;Acc:MGI:1915917] |
| *Trpm6* | transient receptor potential cation channel, subfamily M, member 6 [Source:MGI Symbol;Acc:MGI:2675603] |
| *Trpm7* | transient receptor potential cation channel, subfamily M, member 7 [Source:MGI Symbol;Acc:MGI:1929996] |
| *Trps1* | transcriptional repressor GATA binding 1 [Source:MGI Symbol;Acc:MGI:1927616] |
| *Trpt1* | tRNA phosphotransferase 1 [Source:MGI Symbol;Acc:MGI:1333115] |
| *Trpv1* | transient receptor potential cation channel, subfamily V, member 1 [Source:MGI Symbol;Acc:MGI:1341787] |
| *Trpv2* | transient receptor potential cation channel, subfamily V, member 2 [Source:MGI Symbol;Acc:MGI:1341836] |
| *Trpv3* | transient receptor potential cation channel, subfamily V, member 3 [Source:MGI Symbol;Acc:MGI:2181407] |
| *Trpv4* | transient receptor potential cation channel, subfamily V, member 4 [Source:MGI Symbol;Acc:MGI:1926945] |

| Gene Name | Gene Description |
| --- | --- |
| *Adra1a* | adrenergic receptor, alpha 1a [Source:MGI Symbol;Acc:MGI:104773] |
| *Adra1b* | adrenergic receptor, alpha 1b [Source:MGI Symbol;Acc:MGI:104774] |
| *Adra1d* | adrenergic receptor, alpha 1d [Source:MGI Symbol;Acc:MGI:106673] |
| *Adra2a* | adrenergic receptor, alpha 2a [Source:MGI Symbol;Acc:MGI:87934] |
| *Adra2c* | adrenergic receptor, alpha 2c [Source:MGI Symbol;Acc:MGI:87936] |
| *Adrb1* | adrenergic receptor, beta 1 [Source:MGI Symbol;Acc:MGI:87937] |
| *Adrb2* | adrenergic receptor, beta 2 [Source:MGI Symbol;Acc:MGI:87938] |
| *Adrb3* | adrenergic receptor, beta 3 [Source:MGI Symbol;Acc:MGI:87939] |
| *Agtr1a* | angiotensin II receptor, type 1a [Source:MGI Symbol;Acc:MGI:87964] |
| *Agtr1b* | angiotensin II receptor, type 1b [Source:MGI Symbol;Acc:MGI:87965] |
| *Avpr1a* | arginine vasopressin receptor 1A [Source:MGI Symbol;Acc:MGI:1859216] |
| *Fgf14* | fibroblast growth factor 14 [Source:MGI Symbol;Acc:MGI:109189] |
| *Gucy1a1* | guanylate cyclase 1, soluble, alpha 1 [Source:MGI Symbol;Acc:MGI:1926562] |
| *Gucy1a2* | guanylate cyclase 1, soluble, alpha 2 [Source:MGI Symbol;Acc:MGI:2660877] |
| *Gucy1b1* | guanylate cyclase 1, soluble, beta 1 [Source:MGI Symbol;Acc:MGI:1860604] |
| *Htr2b* | 5-hydroxytryptamine (serotonin) receptor 2B [Source:MGI Symbol;Acc:MGI:109323] |
| *Nedd4l* | neural precursor cell expressed, developmentally down-regulated gene 4-like [Source:MGI Symbol;Acc:MGI:1933754] |
| *Nos1* | nitric oxide synthase 1, neuronal [Source:MGI Symbol;Acc:MGI:97360] |
| *Nos2* | nitric oxide synthase 2, inducible [Source:MGI Symbol;Acc:MGI:97361] |
| *Nos3* | nitric oxide synthase 3, endothelial cell [Source:MGI Symbol;Acc:MGI:97362] |
| *Notch1* | notch 1 [Source:MGI Symbol;Acc:MGI:97363] |
| *Oxtr* | oxytocin receptor [Source:MGI Symbol;Acc:MGI:109147] |
| *Pde3a* | phosphodiesterase 3A, cGMP inhibited [Source:MGI Symbol;Acc:MGI:1860764] |
| *Pik3ap1* | phosphoinositide-3-kinase adaptor protein 1 [Source:MGI Symbol;Acc:MGI:1933177] |
| *Pik3c2a* | phosphatidylinositol-4-phosphate 3-kinase catalytic subunit type 2 alpha [Source:MGI Symbol;Acc:MGI:1203729] |
| *Pik3c2b* | phosphatidylinositol-4-phosphate 3-kinase catalytic subunit type 2 beta [Source:MGI Symbol;Acc:MGI:2685045] |
| *Pik3c3* | phosphatidylinositol 3-kinase catalytic subunit type 3 [Source:MGI Symbol;Acc:MGI:2445019] |
| *Pik3ca* | phosphatidylinositol-4,5-bisphosphate 3-kinase catalytic subunit alpha [Source:MGI Symbol;Acc:MGI:1206581] |
| *Pik3cb* | phosphatidylinositol-4,5-bisphosphate 3-kinase catalytic subunit beta [Source:MGI Symbol;Acc:MGI:1922019] |
| *Pik3cd* | phosphatidylinositol-4,5-bisphosphate 3-kinase catalytic subunit delta [Source:MGI Symbol;Acc:MGI:1098211] |
| *Pik3cg* | phosphatidylinositol-4,5-bisphosphate 3-kinase catalytic subunit gamma [Source:MGI Symbol;Acc:MGI:1353576] |
| *Pik3ip1* | phosphoinositide-3-kinase interacting protein 1 [Source:MGI Symbol;Acc:MGI:1917016] |
| *Pik3r1* | phosphoinositide-3-kinase regulatory subunit 1 [Source:MGI Symbol;Acc:MGI:97583] |
| *Pik3r2* | phosphoinositide-3-kinase regulatory subunit 2 [Source:MGI Symbol;Acc:MGI:1098772] |
| *Pik3r3* | phosphoinositide-3-kinase regulatory subunit 3 [Source:MGI Symbol;Acc:MGI:109277] |
| *Pik3r4* | phosphoinositide-3-kinase regulatory subunit 4 [Source:MGI Symbol;Acc:MGI:1922919] |
| *Pik3r5* | phosphoinositide-3-kinase regulatory subunit 5 [Source:MGI Symbol;Acc:MGI:2443588] |
| *Pik3r6* | phosphoinositide-3-kinase regulatory subunit 6[Source:MGI Symbol;Acc:MGI:2144613] |
| *Pkn1* | protein kinase N1 [Source:MGI Symbol;Acc:MGI:108022] |
| *Pkn2* | protein kinase N2 [Source:MGI Symbol;Acc:MGI:109211] |
| *Pkn3* | protein kinase N3 [Source:MGI Symbol;Acc:MGI:2388285] |
| *Plcb1* | phospholipase C, beta 1 [Source:MGI Symbol;Acc:MGI:97613] |
| *Plcb2* | phospholipase C, beta 2 [Source:MGI Symbol;Acc:MGI:107465] |
| *Plcb3* | phospholipase C, beta 3 [Source:MGI Symbol;Acc:MGI:104778] |
| *Plcb4* | phospholipase C, beta 4 [Source:MGI Symbol;Acc:MGI:107464] |
| *Plcd1* | phospholipase C, delta 1 [Source:MGI Symbol;Acc:MGI:97614] |
| *Plcd3* | phospholipase C, delta 3 [Source:MGI Symbol;Acc:MGI:107451] |
| *Plcd4* | phospholipase C, delta 4 [Source:MGI Symbol;Acc:MGI:107469] |
| *Plce1* | phospholipase C, epsilon 1 [Source:MGI Symbol;Acc:MGI:1921305] |
| *Plcg1* | phospholipase C, gamma 1 [Source:MGI Symbol;Acc:MGI:97615] |
| *Plcg2* | phospholipase C, gamma 2 [Source:MGI Symbol;Acc:MGI:97616] |
| *Plch2* | phospholipase C, eta 2 [Source:MGI Symbol;Acc:MGI:2443078] |
| *Plcl1* | phospholipase C-like 1 [Source:MGI Symbol;Acc:MGI:3036262] |
| *Plcl2* | phospholipase C-like 2 [Source:MGI Symbol;Acc:MGI:1352756] |
| *Plcxd1* | phosphatidylinositol-specific phospholipase C, X domain containing 1 [Source:MGI Symbol;Acc:MGI:2685422] |
| *Plcxd2* | phosphatidylinositol-specific phospholipase C, X domain containing 2 [Source:MGI Symbol;Acc:MGI:3647874] |
| *Prkaca* | protein kinase, cAMP dependent, catalytic, alpha [Source:MGI Symbol;Acc:MGI:97592] |
| *Prkacb* | protein kinase, cAMP dependent, catalytic, beta [Source:MGI Symbol;Acc:MGI:97594] |
| *Prkar1a* | protein kinase, cAMP dependent regulatory, type I, alpha [Source:MGI Symbol;Acc:MGI:104878] |
| *Prkar1b* | protein kinase, cAMP dependent regulatory, type I beta [Source:MGI Symbol;Acc:MGI:97759] |
| *Prkar2a* | protein kinase, cAMP dependent regulatory, type II alpha [Source:MGI Symbol;Acc:MGI:108025] |
| *Prkar2b* | protein kinase, cAMP dependent regulatory, type II beta [Source:MGI Symbol;Acc:MGI:97760] |
| *Prkca* | protein kinase C, alpha [Source:MGI Symbol;Acc:MGI:97595] |
| *Prkcb* | protein kinase C, beta [Source:MGI Symbol;Acc:MGI:97596] |
| *Prkce* | protein kinase C, epsilon [Source:MGI Symbol;Acc:MGI:97599] |
| *Prkcg* | protein kinase C, gamma [Source:MGI Symbol;Acc:MGI:97597] |
| *Prkch* | protein kinase C, eta [Source:MGI Symbol;Acc:MGI:97600] |
| *Prkci* | protein kinase C, iota [Source:MGI Symbol;Acc:MGI:99260] |
| *Prkcz* | protein kinase C, zeta [Source:MGI Symbol;Acc:MGI:97602] |
| *Prkg1* | protein kinase, cGMP-dependent, type I [Source:MGI Symbol;Acc:MGI:108174] |
| *Rapgef1* | Rap guanine nucleotide exchange factor (GEF) 1 [Source:MGI Symbol;Acc:MGI:104580] |
| *Rapgef2* | Rap guanine nucleotide exchange factor (GEF) 2 [Source:MGI Symbol;Acc:MGI:2659071] |
| *Rapgef3* | Rap guanine nucleotide exchange factor (GEF) 3 [Source:MGI Symbol;Acc:MGI:2441741] |
| *Rapgef4* | Rap guanine nucleotide exchange factor (GEF) 4 [Source:MGI Symbol;Acc:MGI:1917723] |
| *S100a1* | S100 calcium binding protein A1 [Source:MGI Symbol;Acc:MGI:1338917] |
| *Sgk1* | serum/glucocorticoid regulated kinase 1 [Source:MGI Symbol;Acc:MGI:1340062] |
| *Slc5a3* | solute carrier family 5 (inositol transporters), member 3 [Source:MGI Symbol;Acc:MGI:1858226] |
| *Src* | Rous sarcoma oncogene [Source:MGI Symbol;Acc:MGI:98397] |
| *Sri* | sorcin [Source:MGI Symbol;Acc:MGI:98419] |
| *Tbxa2r* | thromboxane A2 receptor [Source:MGI Symbol;Acc:MGI:98496] |

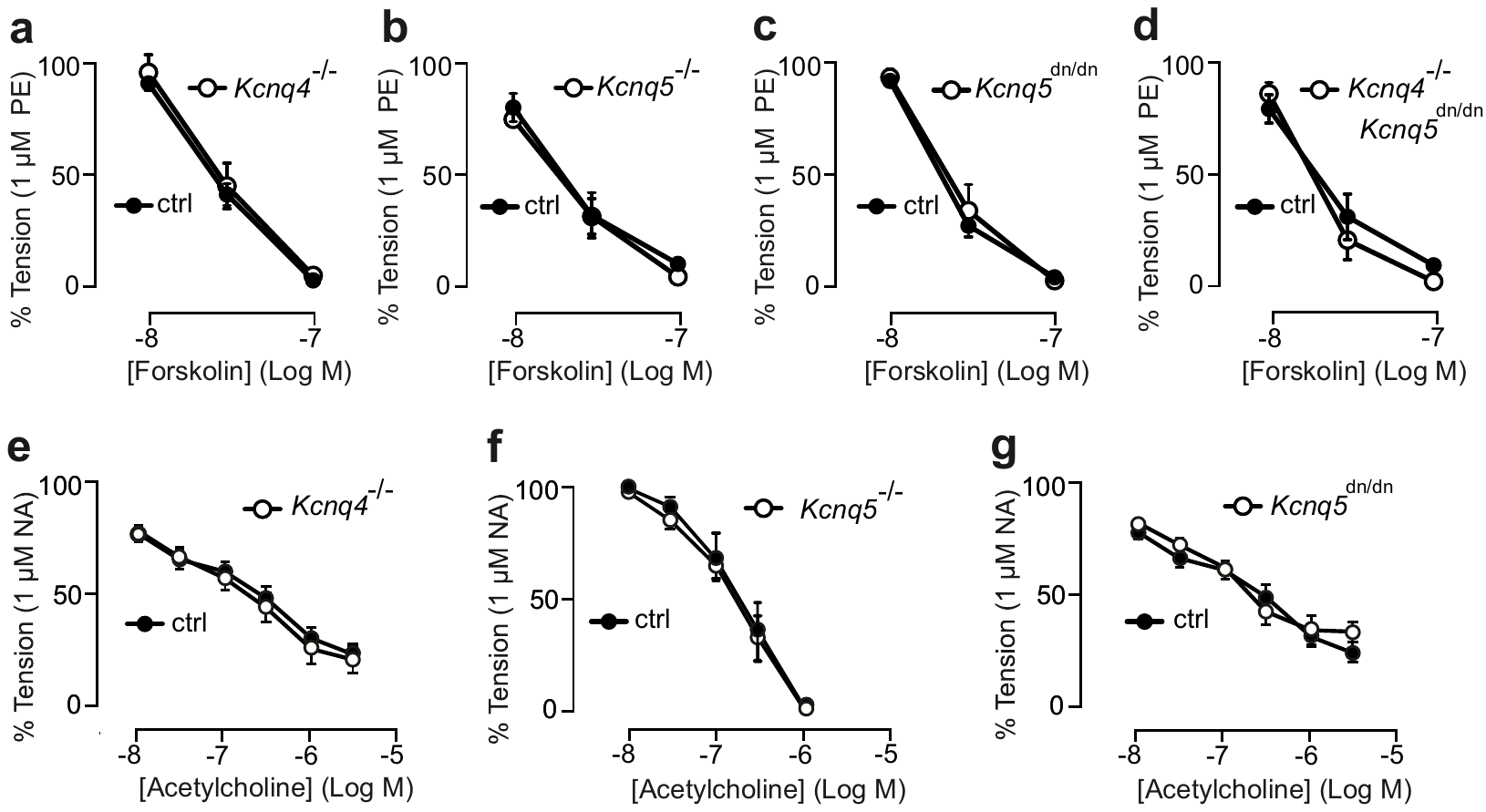

Fig. 5 **Effects of cAMP, NO–cGMP, TRPV4–IK/SK pathways**

Relaxation of (−) PVAT mesenteric arteries by forskolin in *Kcnq4*^−/−^ (**a**), *Kcnq5*^−/−^ (**b**), (*Kcnq5*^dn/dn^ **(c**), *Kcnq4*^−/−^*Kcnq5*^dn/dn^ (**d**), and respective control (*Kcnq4*^+/+^, *Kcnq5*^+/+^, *Kcnq4*^+/+^*/Kcnq5*^+/+^) mice. Relaxation of (−) PVAT mesenteric arteries by acetylcholine in *Kcnq4*^−/−^ (**e**), *Kcnq5*^−/−^ (**f**), or *Kcnq5*^dn/dn^ (**g**) mice. P as determined by two-way ANOVA with Šidák post hoc test. n≥5 arteries from N≥3 mice.

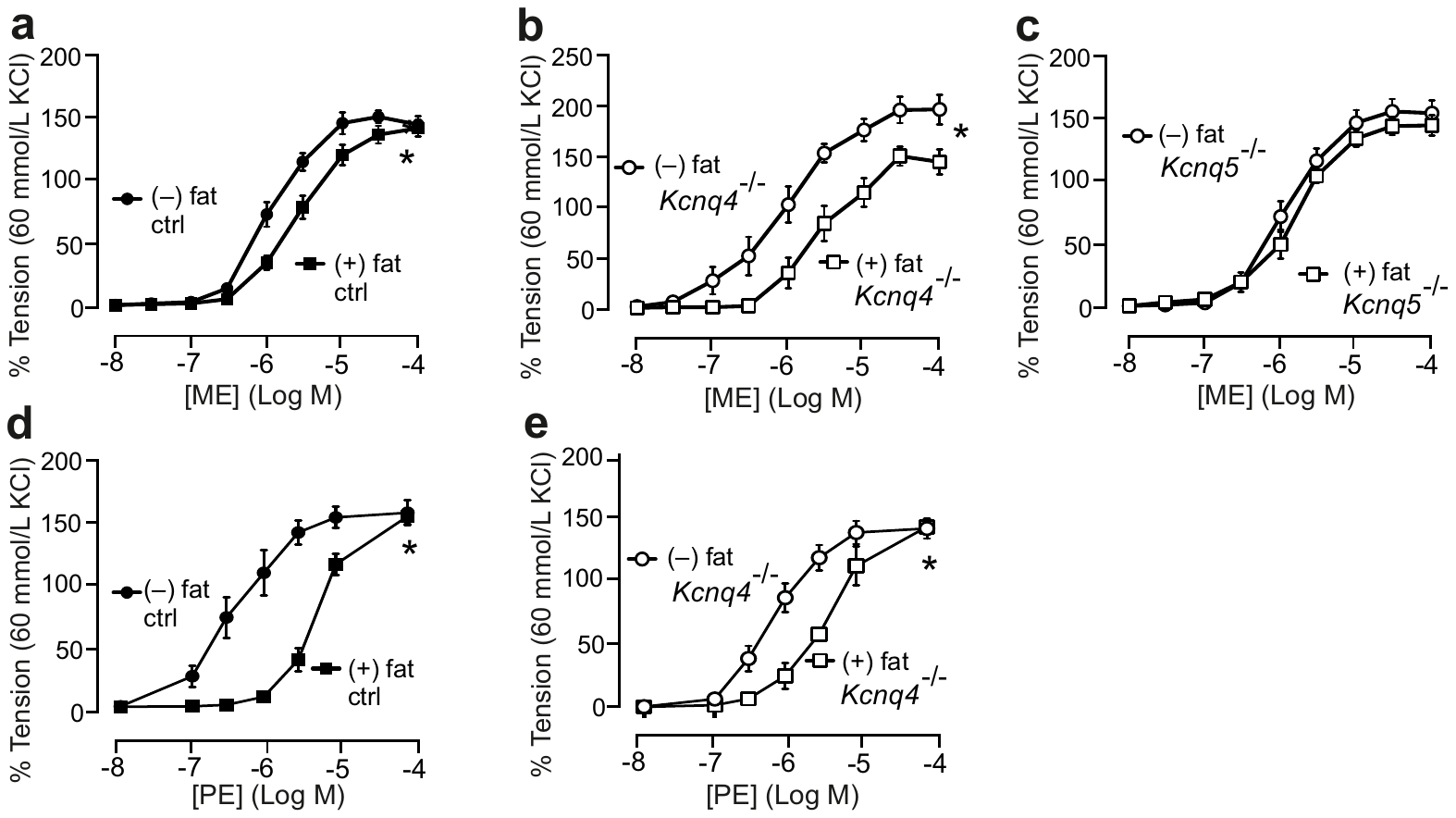

Fig. 6 **KCNQ5 but not KCNQ4 is required for PVAT–mediated control of vascular tone**

Concentration-response relationships for alpha1 adrenoreceptor agonist-induced contractions ME in (+) PVAT or (–) PVAT artery rings isolated from control (**a**), *Kcnq4*^−/−^ (**b**), *Kcnq5*^−/−^ (**c**), and PE in control (**d**) and *Kcnq4*^−/−^ (**e**) mice. Tension is expressed as a percentage of 60 mM KCl-induced contractions. n≥6 arteries from N≥3 mice. Data are mean ± s.e.m. *P as determined by two-way ANOVA with Šidák post hoc test.

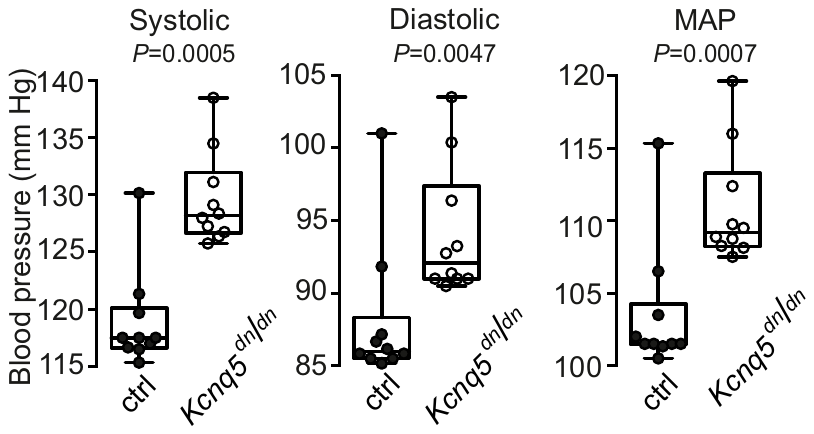
Fig. 7 **KCNQ5 deficiency increases blood pressure**

Systolic, diastolic and mean arterial blood pressures (MAP) over 10 days in control *Kcnq5*^+/+^ (n=6) or *Kcnq5*^dn/dn^ mice (n=8 mice per group). Box plots show median and IQR, whiskers are max and min values. *P* as determined by two-tailed Mann–Whitney *U* test.

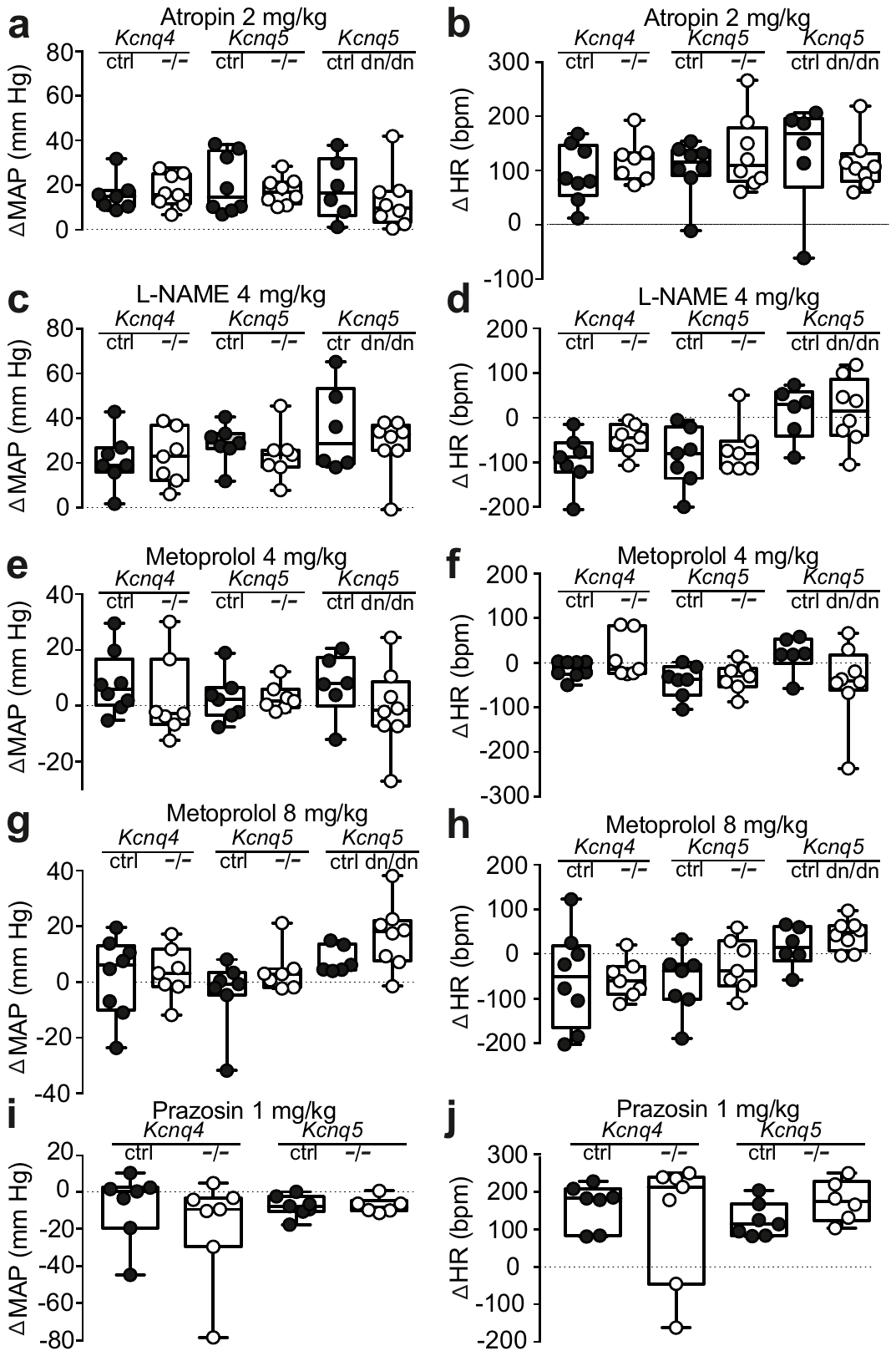

Fig. 8 **Baroreceptor reflex is not changed in KCNQ4 and KCNQ5 deficient animals**

Baroreflex sensitivity in *Kcnq4*^−/−^, *Kcnq5*^−/−^, *Kcnq5*^dn/dn^ and respective control mice. Changes of mean arterial blood pressure (MAP) and heart rate (HR) upon atropine (2 mg/kg) (**a**, **b)**, L-NAME (4 mg/kg) (**c**, **d**), metoprolol (4 mg/kg) (**e**, **f**), metoprolol (8 mg/kg) (**g**, **h**), and prazosin (1 mg/kg) (**i**, **j**) application versus baseline. P as determined by two-way ANOVA with Šidák post hoc test.

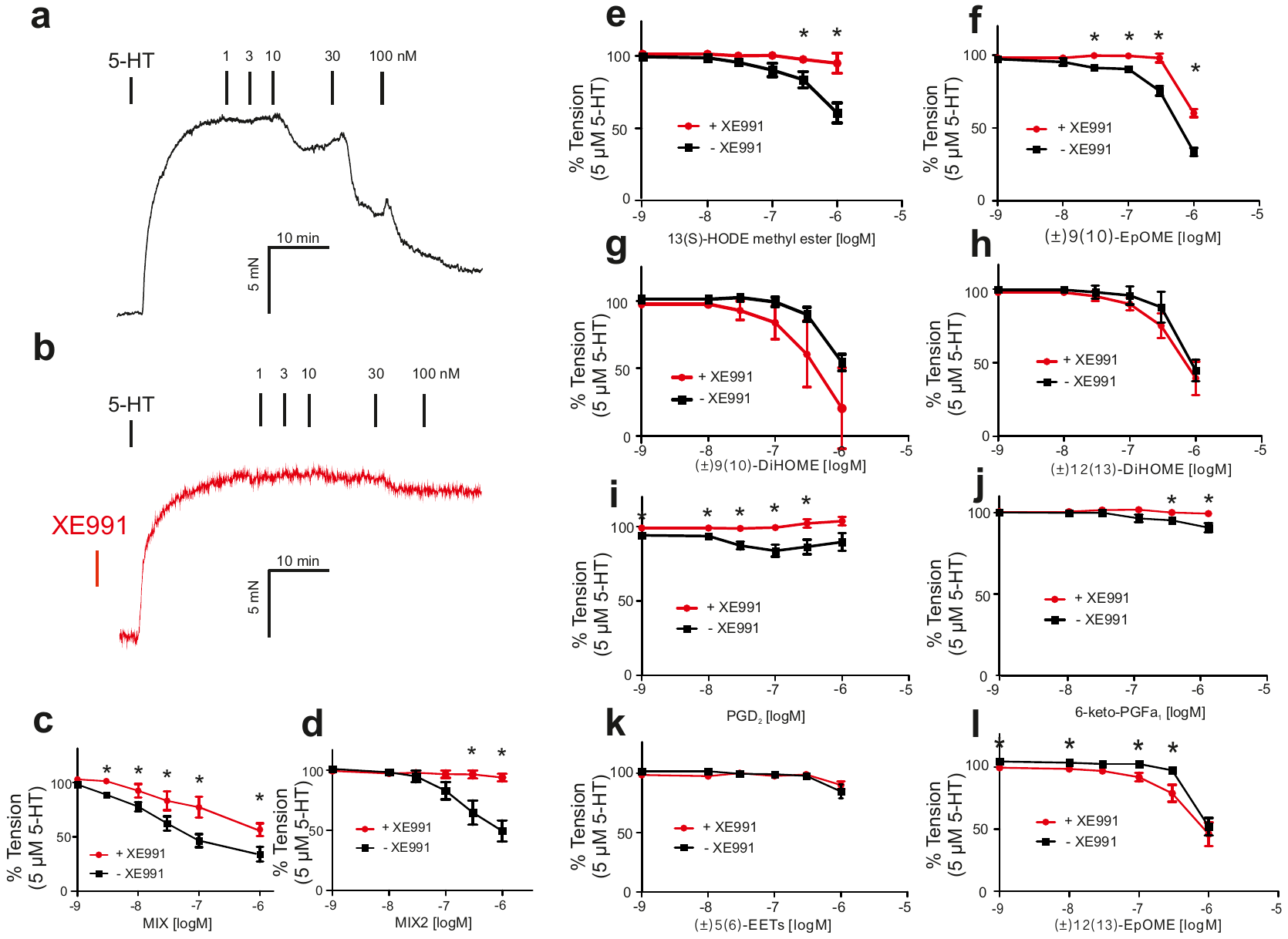

Fig. 9 **Mixture of detected oxylipins relaxes rat aorta stronger than individual substances**

Relaxation of (−) PVAT aortic rings by lipid mediators released from PVAT. Original recordings demonstrating the relaxant effect of oxylipins mixture in the absence (black traces, (**a**)) and presence (red traces, (**b**)) of pan-K_V_7 inhibitor (30 µM XE991). Concentration-response relationships for oxylipins mixtures MIX (**c**), MIX2 (**d**), and individual substances (13(S)-HODE Methyl ester (**e**), (±)9(10)-EpOME (**f**), (±)9(10)-DiHOME (**g**), (±)12(13)-DiHOME (**h**), PGD2 (**i**), 6-keto-PGFa1 (**j**), (±)5(6)-EET (**k**), (±)12(13)-EpOME (**l**). MIX and MIX2 was applied as 1 : 1 : 1 : 1 : 1 : 1 : 1 : 1 and 1 : 0.080 : 0.0950 : 0.351 : 0.123 : 0.362 : 0.017 : 0.001 ratios, respectively. N≥6 rats. *, p<0.05, as determined by two-way ANOVA with Šidák post hoc test.

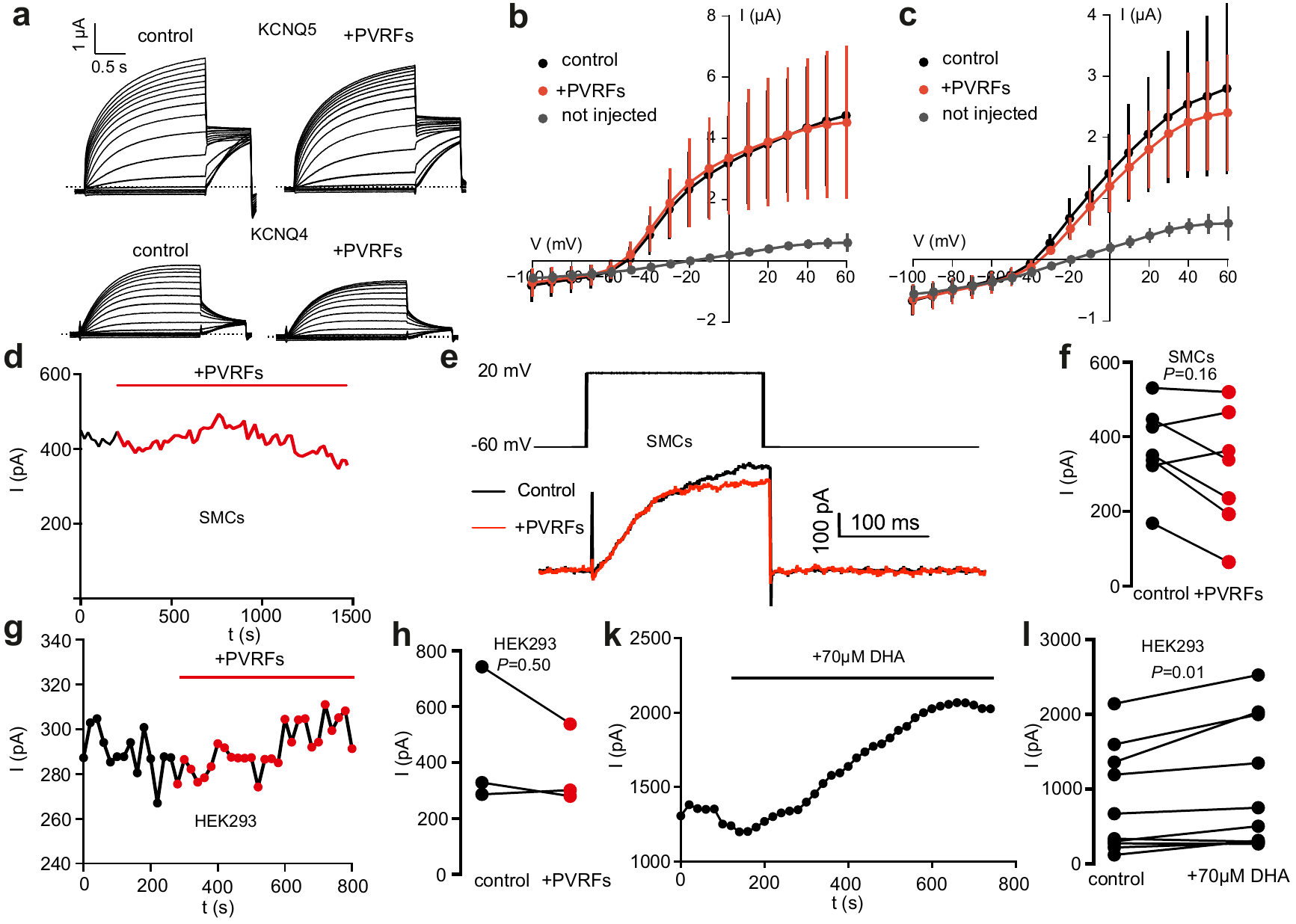

Fig. 10 **PVRFs modulate KCNQ channel activity indirectly**

Averaged current traces (n = 5) from oocytes injected with KCNQ5 or KCNQ4 (**a**) cRNA, with background currents subtracted (estimated from n = 4 non-injected oocytes) before (control) and after (+PVRFs) application of PVRFs.

Current-voltage relationship measured in oocytes injected with KCNQ5 (**b**) or KCNQ4 (**c**) cRNA or uninjected oocytes (gray curve in both **b** and **c**) before (shown in black) and after (in red) application of the PVRFs at the end of each 2-s voltage step. Data is shown as mean ± S.E.M. Representative time course of peak current amplitudes in the absence and presence PVRFs in freshly isolated SMCs (**d, e**). Effect of PVRFs on currents attributed to KCNQ channels in SMCs cells (**f**). Representative time course of peak current amplitudes in the absence and presence PVRFs in HEK293 cells (**g**). Effect of PVRFs on currents attributed to heterologously expressed KCNQ5 channels in HEK293 cells (**h**). Test pulses (200 ms) to +20 mV from -60 mV holding potential; test pulse frequency, 1/20 s. HEK293 cells were transfected using Roti®-Fect according to the vendor’s instructions. Effects of DHA on K_v_ currents in HEK293 cells heterologously expressing KCNQ5 channels (**k**). Representative time course of peak current amplitudes in the absence and presence of 70 μM DHA. Voltage clamp protocol included test steps (200 ms) to 20 mV. Holding potential was -60 mV; test pulse frequency, 1/20 s. HEK293 cell was transfected using Roti®-Fect according to the vendor’s instructions. Effects of DHA on K_v_ currents in presence of 70 µM docosahexaenoic acid (DHA) (**l**).

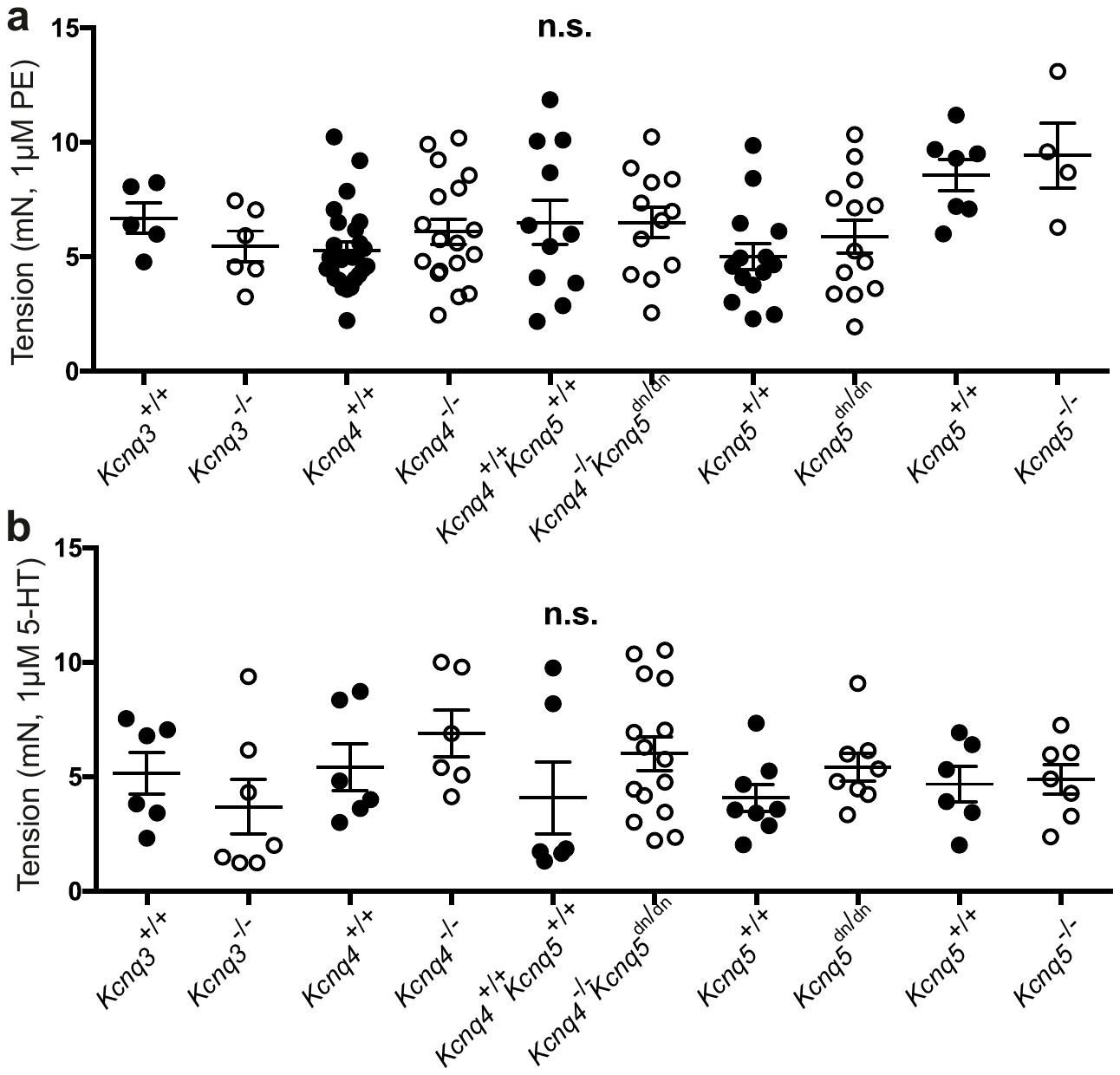

Fig. 11 **Alpha-1 adrenoreceptor (a) and 5-HT (b) contraction was similar across all mice used in the study.**

**Table 1.**

| Compound | p.adj Control vs ME | p.adj Control vs 5-HT |
| --- | --- | --- |
| 12,13-DiHOME | 0.82 | 0.84 |
| 9,10-DiHOME | 0.82 | 0.84 |
| 9-oxo-ODE | 1.00 | 0.95 |
| trans-EKODE | 0.72 | 0.62 |
| 9-HOTrE | 0.82 | 0.53 |
| 5-HETE | 0.72 | 1.00 |
| 17,18-DiHETE | 0.95 | 0.62 |
| 14,15-DHET | 1.00 | 1.00 |
| 5,6-EET | 0.82 | 0.75 |
| PGF1a 6-keto | 0.72 | 0.53 |
| PGD2 | 0.82 | 0.53 |
| LXB4 | 0.72 | 0.53 |
